# Distinct *Wolbachia* localization patterns in oocytes of diverse host species reveal multiple strategies of maternal transmission

**DOI:** 10.1101/2022.10.28.514302

**Authors:** Yonah A. Radousky, Michael T.J. Hague, Sommer Fowler, Eliza Paneru, Adan Codina, Cecilia Rugamas, Grant Hartzog, Brandon S. Cooper, William Sullivan

## Abstract

A broad array of endosymbionts radiate through host populations via vertical transmission, yet much remains unknown concerning the cellular basis, diversity and routes underlying this transmission strategy. Here we address these issues, by examining the cellular distributions of *Wolbachia* strains that diverged up to 50 million years ago in the oocytes of 18 divergent *Drosophila* species. This analysis revealed three *Wolbachia* distribution patterns: 1) a tight clustering at the posterior pole plasm (the site of germline formation); 2) a concentration at the posterior pole plasm, but with a significant bacteria population distributed throughout the oocyte; 3) and a distribution throughout the oocyte, with none or very few located at the posterior pole plasm. Examination of this latter class indicates *Wolbachia* accesses the posterior pole plasm during the interval between late oogenesis and the blastoderm formation. We also find that one *Wolbachia* strain in this class concentrates in the posterior somatic follicle cells that encompass the pole plasm of the developing oocyte. In contrast, strains in which *Wolbachia* concentrate at the posterior pole plasm generally exhibit no or few *Wolbachia* in the follicle cells associated with the pole plasm. Taken together, these studies suggest that for some *Drosophila* species, *Wolbachia* invade the germline from neighboring somatic follicle cells. Phylogenomic analysis indicates that closely related *Wolbachia* strains tend to exhibit similar patterns of posterior localization, suggesting that specific localization strategies are a function of *Wolbachia*-associated factors. Previous studies revealed that endosymbionts rely on *one* of two distinct routes of vertical transmission: continuous maintenance in the germline (germline-to-germline) or a more circuitous route via the soma (germline-to-soma-to-germline). Here we provide compelling evidence that *Wolbachia* strains infecting *Drosophila* species maintain the diverse arrays of cellular mechanisms necessary for *both* of these distinct transmission routes. This characteristic may account for its ability to infect and spread globally through a vast range of host insect species.

## INTRODUCTION

A large class of endosymbionts rely on efficient vertical transmission to ensure their maintenance and spread through host populations. Many insect endosymbionts achieve this by a strict association with the host germline through successive generations. That is, during oogenesis, endosymbionts target the extreme posterior of the oocyte, the site of germline formation. In contrast to this strict germline-to-germline route of maternal transmission, other endosymbionts occupy the germline via invasion from neighboring somatic cells (Russell et al., 2019). Occupation of the germline via a somatic route requires that the endosymbiont has the capacity to undergo cell-to-cell transmission.

Among all known endosymbionts, maternally transmitted *Wolbachia pipientis* is the most common, infecting 30 to 60% of all insect species (Hilgenboecker et al., 2008; Zug and Hammerstein, 2012; Weinert et al., 2015). Hosts acquire *Wolbachia* from other species via horizontal (non-sexual) transfer or from sister species via introgressive transfer (O’Neill et al., 1992; Raychoudhury et al., 2009; Gerth and Bleidorn, 2016; Conner et al., 2017; Turelli et al., 2018; Cooper et al., 2019). Following establishment, maternal transmission rates in conjunction with *Wolbachia* effects on host fitness and reproduction determine *Wolbachia* spread and prevalence in host populations (Hoffmann et al., 1990; Turelli and Hoffmann, 1995; Cooper et al., 2017; Cogni et al., 2021).

While sometimes imperfect, the success and prevalence of *Wolbachia* ultimately depends on vertical transmission through the female (Hoffmann et al., 1990; Turelli and Hoffmann, 1995; Carrington et al., 2011; Hague et al., 2022). The ability of *Wolbachia* to establish and persist in diverse insect species indicates that it has evolved to usurp highly conserved host processes and components. For example, studies of *w*Mel *Wolbachia* in model organism *D. melanogaster* suggest that *Wolbachia* engage conserved host motor proteins Dynein and Kinesin in order to target the extreme posterior of the oocyte, the site of germline formation (Ferree et al., 2005; Serbus and Sullivan, 2007; Russell et al., 2018).

To study the vertical transmission of *Wolbachia* in *Drosophila* we take advantage of the wealth of knowledge regarding the molecular and cell biology of *Drosophila* oogenesis (for a review, see Bastock and St Johnston, 2008). *Drosophila* oocytes contain two clusters of ovarioles, each containing two to three germline stem cells and various stages of developing oocytes. The stem cells undergo asymmetric mitotic division in which one daughter cell undergoes self-renewal and the other differentiates into a cytoblast. The cytoblast undergoes four rounds of mitotic divisions resulting in 16 interconnected cells, one of which becomes the oocyte nucleus while the others differentiate into polyploid nurse cells. The developing oocyte is encapsulated by a monolayer of somatically derived follicle cells resulting in the formation of discrete egg chambers as the maturing oocyte progresses through the ovariole. During the late stages of oogenesis, the nurse cell cytoplasm is expelled into the oocyte, resulting in a rapid doubling of the oocyte volume and concomitant shrinking of nurse cell volume (Theurkauf and Hazelrigg, 1998; Bastock and St Johnston, 2008). This is followed by posterior pole plasm formation and establishment of the germline (Bilinski et al., 2017).

*Wolbachia* distribution and localization in insect oocytes and early embryos has been well characterized in *w*Mel-infected *D. melaongaster* (Kose and Karr, 1995; Veneti et al., 2004; Ferree et al., 2005; Serbus et al., 2008; Ramalho et al., 2018; Russell et al., 2018; Guo et al., 2019). Early in oogenesis, *w*Mel *Wolbachia* are evenly distributed throughout all the cells in the chamber. At this stage, microtubules emanate from the oocyte into the nurse cells (Theurkauf, 1994; Grieder et al., 2000). By associating with the minus-end directed microtubule motor protein Dynein, *w*Mel *Wolbachia* are transported from the nurse cells and concentrate at the anterior end of the oocyte (Ferree et al., 2005). As the oocyte matures, microtubules undergo a dramatic reorganization such that they originate from the oocyte cortex with their plus-ends oriented toward the posterior pole (Theurkauf et al., 1992; Steinhauer and Kalderon, 2006). This reorganization is accompanied by a release and dispersal of *w*Mel *Wolbachia* from the anterior pole. Then, *w*Mel *Wolbachia* associate with the plus-end directed motor protein Kinesin for transport to the posterior pole and the site of germline formation (Serbus and Sullivan, 2007). There, *Wolbachia* (*w*Mel) associate with pole plasm determinants maintaining their position at the site of germline formation (Serbus and Sullivan, 2007) in spite of extensive cytoplasmic streaming (Serbus et al., 2005).

Thus, *w*Mel *Wolbachia* location and migration patterns provide insights into the host factors with which *Wolbachia* engages throughout oogenesis. The *D. simulans*-associated *w*Ri strain exhibits a similar anterior distribution in early oogenesis, and then becomes distributed across the entire mature oocyte. However, unlike *w*Mel, it does not concentrate at the posterior pole of the oocyte (Veneti et al., 2004; Serbus and Sullivan, 2007). This suggests *w*Ri engages host Dynein and Kinesin, but not pole plasm determinants. *w*Mel *Wolbachia* strains introgressed into a *D. simulans* host background exhibit a concentration in the pole plasm, indicating the *Wolbachia* genome rather than the host genome drives oocyte localization properties (Poinsot et al., 1998; Veneti et al., 2004; Serbus and Sullivan, 2007).

With the exception of *w*Mel and *w*Ri, little is known of the extent to which distribution patterns in the oocyte and transmission routes vary among diverse *Wolbachia* strains and host species. Here, we fill this important knowledge gap, by examining 18 *Drosophila* host species infected with *Wolbachia* strains that diverged up to 50 million years ago. These studies also provide a direct read-out of *Wolbachia* and host motor protein/pole plasm interactions that are highly conserved and those that vary among species. This work complements and extends previous analyses revealing *Wolbachia* strain-dependent localization patterns in the *Drosophila* oocyte and embryo (Veneti *et al*., 2004; Toomey et al., 2013).

We discovered that *Wolbachia* distribution patterns in the *Drosophila* oocytes fall into three categories: (1) the vast majority of *Wolbachia* clusters at the posterior pole; (2) a concentration at the posterior pole, but with a large portion of *Wolbachia* distributed throughout the entire oocyte; and (3) *Wolbachia* distributed throughout oocyte, with none or very few located at the posterior pole. This latter class is particularly interesting as it suggests *Wolbachia* likely enters the oocyte via a somatic route for transmission to the next generation. We also provide evidence that neighboring somatic follicle cells are the likely source of *Wolbachia* entering the germline. Together, these findings indicate that *Wolbachia* are capable of achieving efficient vertical transmission either via strict maintenance in the germline from one generation to the next or via invasion of the germline through neighboring somatic cells. Thus, *Wolbachia* must maintain the diverse arrays of cellular mechanisms necessary for *both* of these distinct transmission routes. We identify candidate surface proteins that may be responsible for these diverse localization patterns in host oocytes.

## RESULTS

### *Wolbachia* navigates the developing *Drosophila* oocyte in distinct stages

As described above, *w*Mel *Wolbachia* migration and distribution throughout oogenesis in *D. melanogaster* has been well documented (Kose and Karr, 1995; Veneti et al., 2004; Serbus and Sullivan, 2007; Serbus et al., 2008; Ramalho et al., 2018; Russell et al., 2018; Guo et al., 2019). As a reference for our analysis of *Wolbachia (w*Mel) oocyte distributions in a range of *Drosophila* species, the key distribution patterns of *w*Mel throughout *D. melanogaster* oocytes are depicted in **Figure 1**.

**Figure 1.**
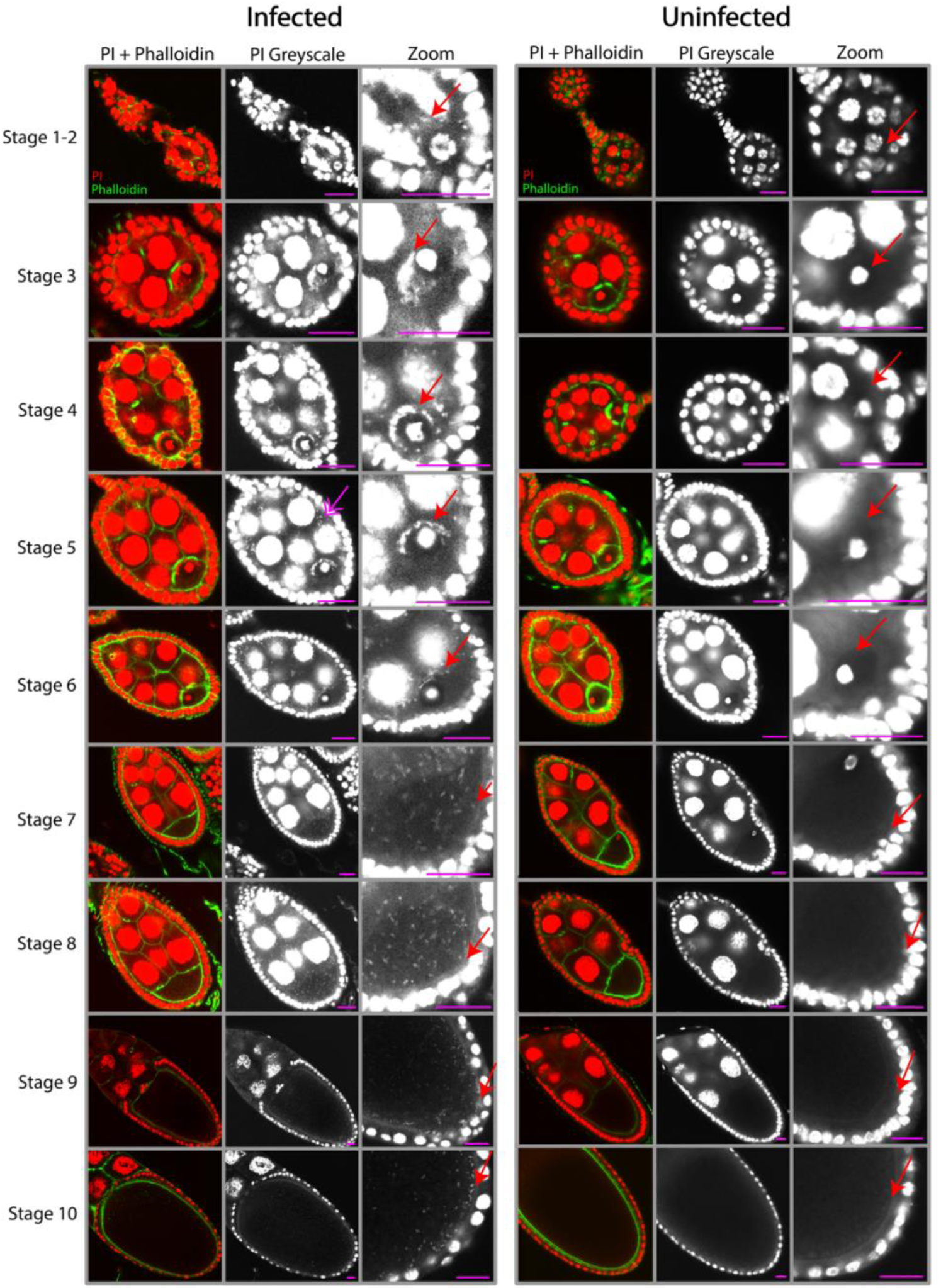
Confocal micrographs of *D. melanogaster* (*w*Mel) oocytes tracked through early oogenesis. Samples are stained with propidium iodide (PI) for DNA (red) and phalloidin for actin (green). Representative stages of development (1-10) are shown for *w*Mel infected (left panels) and uninfected (right panels). Grayscale zoomed-in images around the nucleus of the oocyte are shown in the third column, with red arrows highlighting *Wolbachia* in the oocyte. Corresponding arrows in the sixth column show the equivalent area in uninfected oocytes. The magenta double arrow in the stage 5 infected oocyte (second column) highlights *Wolbachia* in the nurse cells. Scale bars all set to 25 uM.

Stage 1 includes the germarium, a structure consisting of the germline stem cells and their cystoblast daughter cells that will differentiate into a mature oocytes. Progression to stage 2, involves the cystoblast undergoing four rounds of mitotic division to produce a syncytium of 16 cells connected by cytoplasmic bridges and surrounded by a layer of somatically derived follicle cells (Bastock and St Johnston, 2008). Propidium Iodine (PI)-stained *Wolbachia* (red) are clearly observed as puncta in the cytoplasm of the infected, but not the uninfected 16 cell cyst (**Figure 1 Stages 1-2, Arrows**). Actin is labeled in green.

During stage 3, the cyst has become polarized with one of the cells differentiating into an oocyte and the others becoming polyploid nurse cells. At this stage, *Wolbachia* rely on Dynein-mediated microtubule transport (Ferree et al., 2005) and are clearly seen in the infected oocyte cytoplasm anteriorly concentrated at two cytoplasmic bridges that connect to the nurse cells (**Figure 1, Stage 3, Arrows**).

During stages 4 and 5, the nurse cell/oocyte complex becomes oblong and is fully encapsulated by follicle cells. *Wolbachia* are clearly observed in both the oocyte and nurse cell cytoplasm (**Figure 1, Stages 4 and 5, Arrows**). The *Wolbachia* in the oocyte are tightly associated with the anterior cortex.

During stage 6, the number of follicle cells increases and they are more densely packed in a monolayer encompassing the oocyte. Here we discovered a previously unrecognized aspect of *Wolbachia* dynamics and localization: the number of anteriorly localized *Wolbachia* is greatly diminished at this stage (**Figure 1, Stage 6, Arrow**). This pattern was consistent across all 10 of the stage 6 oocytes we observed in our analysis (see **Supplemental Figure 1**).

At stage 7, *Wolbachia* are again observed in the oocyte. However, rather than being tightly associated with the anterior cortex, *Wolbachia* are evenly distributed throughout the oocyte (**Figure 1, Stage 7, Arrow**). During stage 7, the microtubules reorient such that their plus-ends extend toward the posterior pole (Cha et al., 2001; Cha et al., 2002; Steinhauer and Kalderon, 2006). Accordingly, the plus-end motor protein Kinesin is required to establish the even distribution of *Wolbachia* throughout the maturing oocyte (Serbus and Sullivan, 2007). Stage 8 heralds the beginning of vitellogenesis, the process of yolk formation and deposition of nutrients into the oocyte (Cummings et al., 1971). *Wolbachia* are near, but not yet located at, the posterior pole (**Figure 1, Stage 8, Arrow**). At stage 9, anterior follicle cell migration occurs (Rørth, 2002). At this stage, *Wolbachia* displays an archlike posterior cortical localization. (**Figure 1, Stage 9, Arrow**). By stage 10, there is a distinct concentration of *Wolbachia* along the posterior cortex, the site of germline formation (**Figure 1, Stage 10, Arrow**). Nurse cell dumping will shortly follow, leading into stage 11-12 with the oocyte rapidly growing in size (Mahajan-Miklos and Cooley, 1994).

The *w*Mel strain in *D. melanogaster* provides an outline of the molecular and cellular mechanisms driving *Wolbachia* vertical transmission. It also raises the question of whether the mechanisms and strategies observed in *D. melanogaster* are conserved across diverse *Wolbachia* strains and *Drosophila* species. To explore this issue, the *w*Mel distribution in *D. melanogaster* serves as the basis for our comparison of 18 other *Wolbachia* strains. Bayesian phylograms depicting phylogenetic relationships among the *Wolbachia* strains and the *Drosophila* species are shown in **Figure 4** and **Supplemental Figure 2**. Our analysis comprised 18 A-group *Wolbachia* strains, including six *w*Mel-like strains (Cooper et al., 2019; Hague et al., 2020a) and five *w*Ri-like strains (Turelli et al., 2018), as well as the B-group *Wolbachia* strain *w*Mau, which diverged from A-group *Wolbachia* up to 50 million years ago (Meany et al., 2019). These *Wolbachia* strains infect 18 divergent *Drosophila* host species spanning three species groups (Suvorov et al., 2022).

### An initial concentration of *Wolbachia* at the oocyte anterior is a conserved feature

As described above, *w*Mel *Wolbachia* enter the oocyte through a complex of ring canals connecting the nurse cells to the oocyte and concentrate at the anterior cortex at stage 3-5. Initially observed in *D. melanogaster*, the number of *Wolbachia* increases dramatically while at the anterior (Ferree et al., 2005). To determine if anterior localization is a conserved aspect of *Wolbachia*’s navigation through the developing oocyte, we first characterized this trait in eight *Wolbachia* strains in their respective hosts: *w*Pan in *D. pandora*, *w*Nik in *D. nikananu*, *w*Cha in *D. chauvacae*, *w*Ha in *D. simulans*, *w*Tsa in D*. tsacasi*, *w*Boc in *D. bocki*, *w*Ana in *D. ananassae*, and *w*Mel in *D. melanogaster*. All eight strains exhibited a distinct anterior localization during mid-oogenesis **(Figure 2)**. It is interesting that this localization occurs during stage 5 of oogenesis, which, in *D. melanogaster,* is prior to the localization of all known anterior axis determinants. Thus, *Wolbachia* likely relies on as yet undiscovered anteriorly concentrated factor(s).

**Figure 2.**
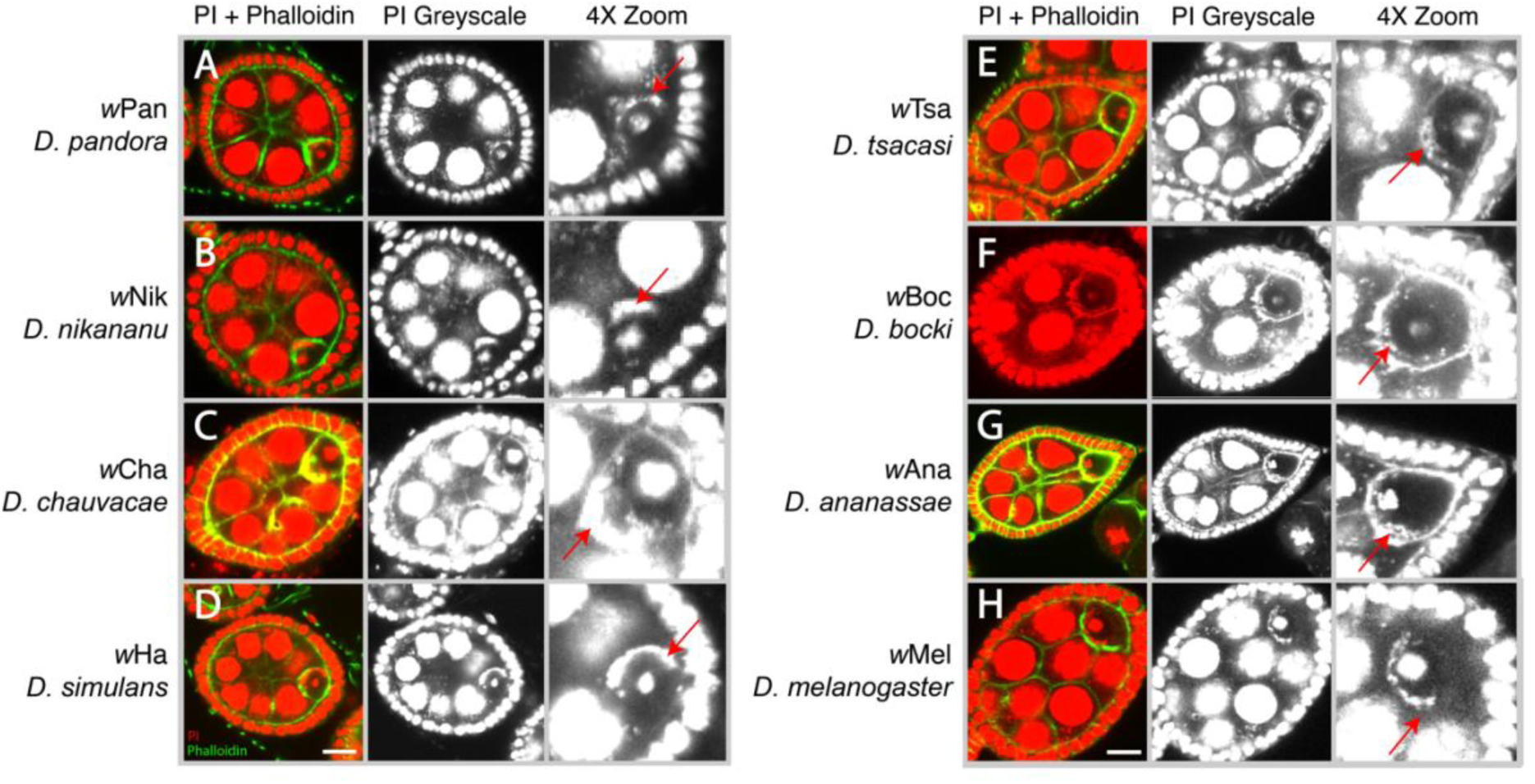
Conserved *Wolbachia* anterior localization during early oogenesis (stages 4-5). *Wolbachia* and host DNA stained with PI (red) and actin with phalloidin (green) shown in the left most column. Greyscale of PI channel shown in middle. 4X zoom-in around the oocyte shown on the right. All of the eight *Wolbachia* strains examined concentrate at the anterior cortex of the oocyte at this stage. Red arrows highlight *Wolbachia*’s anterior localization positioning for (A) *w*Pan in *D. pandora*, (B) *w*Nik in *D. nikananu*, (C) *w*Cha in *D. chauvacae*, (D) *w*Ha in *D. simulans*, (E) *w*Tsa in D*. tsacasi*, (F) *w*Boc in *D. bocki*, (G) *w*Ana in *D. ananassae*, and (H) *w*Mel in *D. melanogaster*. Oocytes are approximately 50-75 uM in size.

### *Wolbachia* exhibits three distinct distributions with respect to posterior localization in the mature *Drosophila* oocyte

Concomitant with microtubule rearrangement during stages 7-8, *w*Mel *Wolbachia* are released from the oocyte anterior resulting in the dispersal of the bacteria throughout the entire length of the oocyte with a fraction concentrating at the posterior pole (Ferree et al., 2005). The posterior dispersal of *Wolbachia* requires the plus-end motor protein Kinesin (Serbus and Sullivan, 2007). Maintenance of those *Wolbachia* that reach the posterior pole rely on a stable association with key pole plasm components (Serbus et al., 2011).

We next examined the oocytes of 18 *Drosophila* species infected with 19 diverse *Wolbachia* strains (*N* = 72 total oocytes) to test the hypothesis that oocyte posterior distributions vary. Unlike the conserved anterior localization, *Wolbachia* exhibit three distinct patterns of posterior localization in stage 10 oocytes. As shown in **Figure 3** and **Supplemental Figure 3A**, eight *Wolbachia* strains exhibit a localization pattern similar to that observed for *w*Mel in *D. melanogaster* with the bulk of *Wolbachia* evenly distributed throughout the oocyte, and a large fraction concentrated at the posterior pole. We refer to this pattern as <u>Posteriorly Localized</u>.

**Figure 3.**
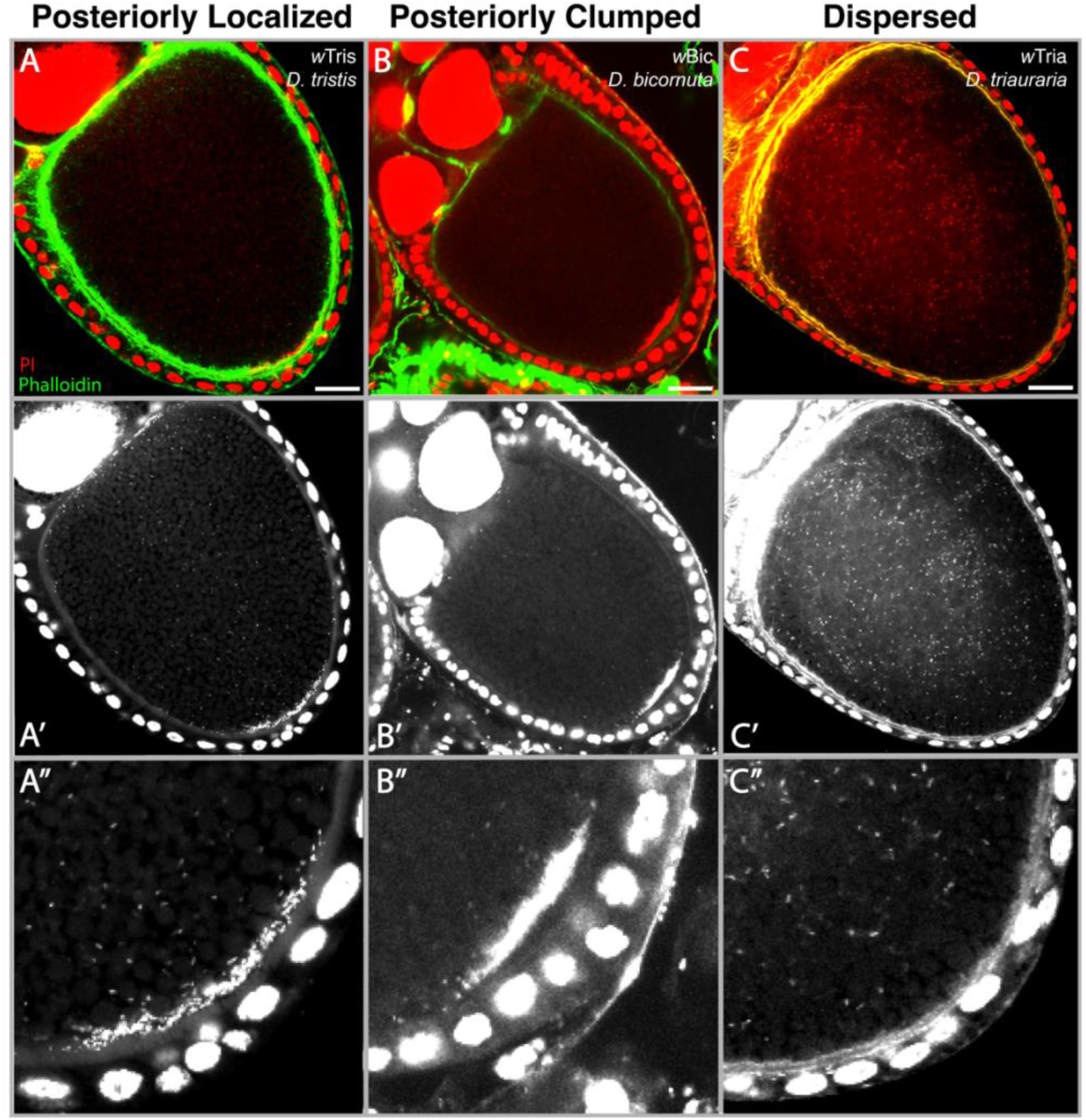
Three distinct patterns of posterior *Wolbachia* localization during late stage oogenesis (A-C). Confocal micrographs of *Drosophila* oocytes DNA-stained with PI (red) and actin-stained with phalloidin (green) show representative examples of (A) Posteriorly Localized *w*Tris in *D. tristis*, (B) Posteriorly Clumped *w*Bic in *D. bicornuta*, and (C) Dispersed localization of *w*Tria in *D. triauraria*. Second row depicts single channel images of PI staining. Third row depicts an enlarged PI-stained image of the posterior region of each oocyte. Scale bars set at 25uM.

Our analysis revealed four *Wolbachia* strains manifested a second pattern in which the vast majority of *Wolbachia* localized to a distinct cluster at the posterior pole and only few *Wolbachia* are distributed throughout the remainder of the oocyte **(Figure 3, Supplemental Figure 3B)**. In contrast to the pattern described above, rather than being distributed in a wide arc along the posterior pole, they are tightly centered at the extreme posterior region of the oocyte (**Figure 3B**). We refer to this pattern as <u>Posteriorly Clumped</u>.

Finally, seven *Wolbachia* strains exhibit a third pattern in which *Wolbachia* are distributed throughout the entire oocyte but there are few to no bacteria concentration at the posterior pole (**Figure 3 and Supplemental Figure 3C**). The lack of the *Wolbachia* at the posterior pole was unexpected as this was thought to be necessary for incorporation into the germline of the next generation. We refer to this pattern as <u>Dispersed</u>.

### Quantification of *Wolbachia* oocyte distribution patterns

We used one-way ANOVAs to test whether the three classes (Posteriorly Localized, Posteriorly Clumped, and Dispersed) differ statistically in quantitative estimates of *Wolbachia* abundance in the whole oocyte, at the posterior region, and at the posterior cortex (**Supplemental Figure 4**). We defined the posterior region as the posterior 12.5% of the oocyte, and the posterior cortex as the narrow cortical region of Vasa expression (Russell et al., 2018), and we then measured total *Wolbachia* abundance in each region as the Corrected Total Cell Fluorescence (CTCF). We found that *Wolbachia* abundance differs significantly among the three classes (one-way ANOVA, *F*_(2,69)_ = 6.2, *P* = 0.003), with Tukey multiple comparisons indicating that the Clumped class has significantly more *Wolbachia* in whole oocytes than the Localized group (*P* = 0.002). The three classes also differed significantly in the amount of *Wolbachia* in the posterior region (*F*_(2,69)_ = 23.09, *P* < 0.001), such that the Dispersed class has significantly fewer *Wolbachia* at the posterior that the Clumped (*P* < 0.001) and Localized (*P* < 0.001) groups. Finally, the three classes also differ in the amount of *Wolbachia* at the posterior cortex (*F*_(2,64)_ = 38.12, *P* < 0.001). Here, multiple comparisons indicate that all three classes significantly differ in levels of *Wolbachia* at the posterior cortex: the Clumped class has the most *Wolbachia*, followed by Localized, and then Dispersed (**Supplemental Figure 4**). Mean estimates of *Wolbachia* abundance (CTCF) for each *Wolbachia* strain and host species are shown in **Table 1**.

**Table 1.** Qualitative and quantitative quantification of cellular Wolbachia abundance in stage 10 oocytes. The number of confocal images (N) used to generate means is shown for each Wolbachia-infected host species. Wolbachia abundance was measured in each region of the oocyte as corrected total cell fluorescence (CTCF).

| *Wolbachia* strain | Host species | Localization class | $N$ | Whole oocyte mean CTCF | Posterior mean CTCF | Posterior cortex mean CTCF |
| --- | --- | --- | --- | --- | --- | --- |
| wRec | *D. recens* | Clumped | 3 | 482174 | 300753 | 58453 |
| wTris | *D. tristis* | Localized | 4 | 223048 | 129013 | 105641 |
| wSeg | *D. seguyi* | Dispersed | 5 | 58552 | 11402 | 557 |
| wCha | *D. chauvacaе* | Localized | 5 | 108588 | 42636 | 9786 |
| wBurl | *D. burlai* | Localized | 4 | 72806 | 53754 | 36944 |
| wMel | *D. melanogaster* | Localized | 2 | 191451 | 136174 | 92341 |
| wBic | *D. bicornuta* | Clumped | 3 | 90114 | 83621 | 54900 |
| wRi | *D. simulans* | Dispersed | 5 | 333505 | 90664 | 20494 |
| wPan | *D. pandora* | Localized | 3 | 45423 | 6486 | 1890 |
| wAna | *D. ananassae* | Localized | 4 | 41578 | 16843 | 12927 |
| wTria | *D. triauraria* | Dispersed | 2 | 487897 | 1809 | 345 |
| wAura | *D. auraria* | Dispersed | 5 | 266478 | 3699 | 645 |
| wHa | *D. simulans* | Dispersed | 5 | 82941 | 3488 | 807 |
| wBai | *D. baimaii* | Dispersed | 2 | 11359 | 0 | 0 |
| wBoc | *D. bocki* | Clumped | 6 | 137664 | 98339 | 82408 |
| wLeo | *D. leontia* | Clumped | 4 | 997045 | 685260 | 713918 |
| wTsa | *D. tsacasi* | Localized | 4 | 61314 | 36478 | 20620 |
| wNik | *D. nikananu* | Localized | 3 | 44764 | 27099 | 13014 |
| wMau | *D. mauritiana* | Dispersed | 3 | 69362 | 799 | 322 |

### *Wolbachia* abundance in the oocyte posterior has strong phylogenetic signal

We hypothesized that the diversity of *Wolbachia* localization patterns may be due to *Wolbachia*-associated factors, implying that closely related *Wolbachia* strains would exhibit similar localization patterns in host oocytes. We used the phylogram describing the evolutionary relationships among *Wolbachia* strains to test whether patterns of *Wolbachia* localization exhibit a phylogenetic signal using Pagel’s λ (Pagel, 1999; Hague et al., 2020b; Hague et al., 2021) (**Figure 4**). A λ value of 1 would be consistent with trait evolution that entirely agrees with the *Wolbachia* phylogeny (i.e., strong phylogenetic signal supporting a role of *Wolbachia* factors), whereas a value of 0 would indicate that trait evolution occurs independently of phylogenetic relationships (Pagel, 1999; Freckleton et al., 2002). Most notably, we found that *Wolbachia* abundance (CTCF) in the posterior region exhibits a strong, significant phylogenetic signal (λ = 0.982 [0, 0.998], *P* = 0.021). *Wolbachia* abundance at the posterior cortex also has a high, but non-significant λ value (λ = 0.959 [0, 0.995], *P* = 0.683). Here, the large confidence intervals and our simulations suggest that a larger *Wolbachia* phylogeny (*N* = 25, 50 *Wolbachia* strains) is required to detect a significant signal of λ > 0 (**Supplemental Figure 5**). These results highlight differences in patterns of *Wolbachia* localization between the *w*Mel- and *w*Ri-like *Wolbachia* clades (**Figure 3**). Closely related *w*Mel-like *Wolbachia* tend to occur at a higher abundance in the posterior region of oocytes, whereas the *w*Ri-like strains are generally more dispersed. We found no evidence that *Wolbachia* abundance in the whole oocyte exhibits a phylogenetic signal (λ < 0.001 [0, 0.753], *P* = 1).

**Figure 4.**
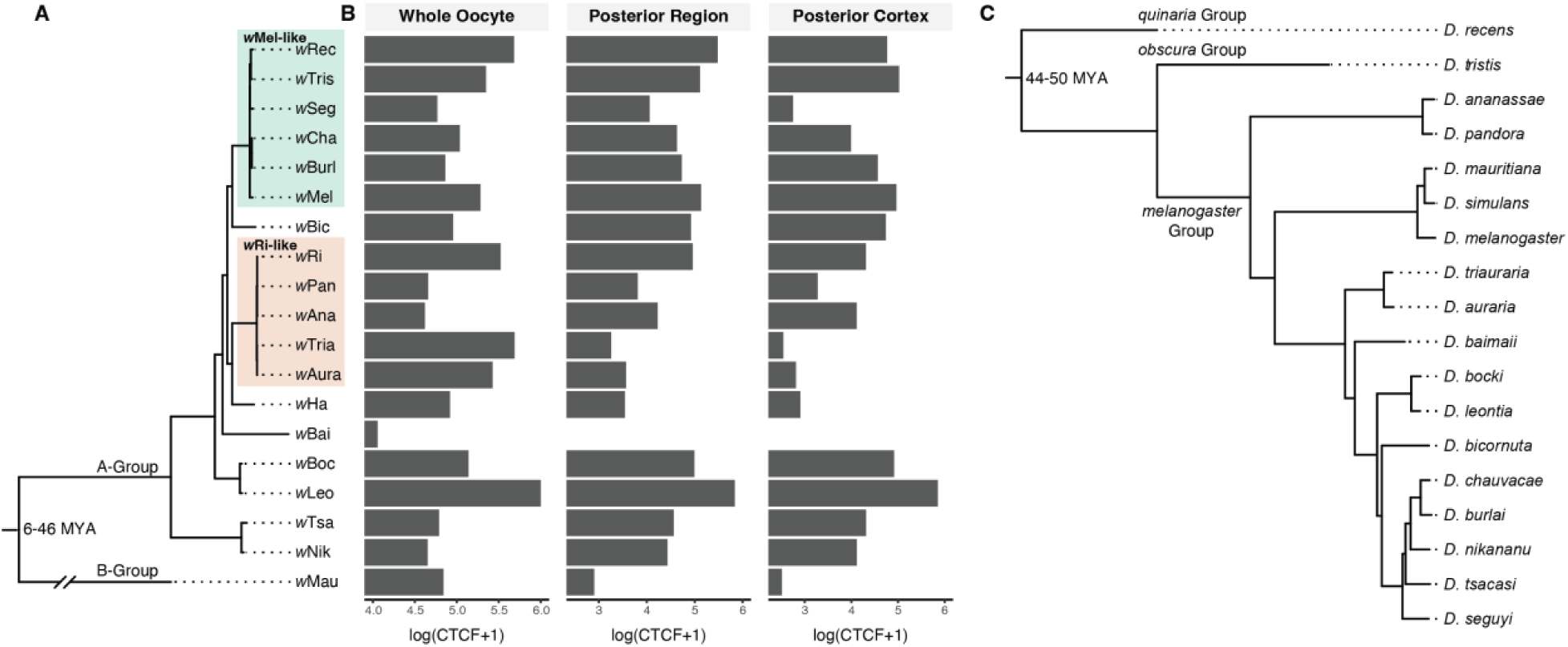
*Wolbachia* abundance in the posterior region of host oocytes exhibits phylogenetic signal on the *Wolbachia* phylogeny. (A) Estimated Bayesian phylogram for the 19 A- and B-group *Wolbachia* strains included in tests for phylogenetic signal. (B) Mean estimates of *Wolbachia* abundance (log-transformed CTCF) in the whole oocyte, posterior region, and the posterior cortex. *Wolbachia* abundance in the posterior region has significant phylogenetic signal (λ = 0.982 [0, 0.998], *P* = 0.021), suggesting *Wolbachia*-associated factors determine *Wolbachia* abundance in the oocyte posterior. (C) Estimated Bayesian phylogram of *Drosophila* host species using 20 conserved single-copy genes. *Wolbachia* and host divergence times in millions of years (MYA) are reproduced from Meany et al. (2019) and Suvorov et al. (2022), respectively. The *Wolbachia* and host tree are generally discordant (see **Supplemental Figure 2**), as expected with frequent horizontal *Wolbachia* acquisition (Turelli et al. 2018).

The correlation between *Wolbachia* localization in oocytes and phylogenetic divergence implies that factors in the *Wolbachia* genome determine *Wolbachia* abundance at the oocyte posterior. Because these distributions likely involve direct interactions between *Wolbachia* and the host cytoplasm, we focused on six major *Wolbachia* surface proteins: the three *wsp* paralogs (WD_1063/*wsp*, WD_0009/*wspB*, WD_0489/*wspC*), as well as WD_0501, WD_1041, WD_1085. We tested whether sequence divergence at any of these candidate loci predicts *Wolbachia* localization patterns. Specifically, we tested whether gene trees of the *Wolbachia* surface proteins (**Supplemental Figure 6**) exhibit a phylogenetic signal, using the methods described above. One surface protein (WD_1085) stood out with especially strong evidence of a phylogenetic signal, with significant departures from λ = 0 for *Wolbachia* abundance at the posterior region (λ = 0.974 [0, 0.998], *P* = 0.001) and the posterior cortex (λ = 0.942 [0, 0.994], *P* = 0.028). In addition, *Wolbachia* abundance at the posterior region also exhibited significant phylogenetic signal on the gene tree of WD_0501 (λ = 0.992 [0.521, 1], *P* = 0.022). All tests for phylogenetic signal using *Wolbachia* gene trees are summarized in **Supplemental Table 1**.

Motivated by previous successful analysis of *Wolbachia* protein function through ectopic expression in yeast, we expressed codon-optimized forms of WD_1085 and WD_0501 in yeast using a Galactose-inducible promoter (Sheehan et al., 2016; Rice et al., 2017). Under normal growth conditions, ectopic expression of both genes inhibited growth, with WD_1085 exhibiting a more pronounced inhibition (**Supplemental Figure 7**). Because *Wolbachia* exhibits a close association with microtubules, we also assayed whether ectopic expression of these genes enhanced the growth defects in situations in which the microtubule spindle assembly checkpoint was compromised by a mad1 mutation (Luo et al., 2018), or when the microtubules were compromised directly through the addition of benomyl. As indicated in **Supplemental Figure 7**, ectopic expression of WD_1085 significantly inhibited growth when both the microtubules and spindle checkpoint were compromised. These results suggest that ectopic WD_1085 expression perturbs microtubule function.

We recently found that the surface protein WD_0009 (*wspB*) is pseudogenized due to a premature stop codon in a tropical *w*Mel variant sampled from Australia, which seems to influence *w*Mel abundance in stage 10 oocytes when hosts are reared in the cold (Hague et al., 2022). Here, we did not find that the *wspB* gene tree exhibits phylogenetic signal related to *Wolbachia* oocyte abundance (**Supplemental Table 1**); however, we note that *wspB* appears to be psuedogenized in a number of other *Wolbachia* strains in addition to tropical *w*Mel. We found derived deletions of varying length in the *wspB* sequences of *w*Ha and *w*Bai beginning at nucleotide position 256, both of which produce frame shifts that generate multiple downstream stop codons (**Supplemental Figure 8**). In addition, *w*Boc and *w*Leo share a large insertion starting at bp position 351. Because the insertions contain a contig break, we were unable to resolve the full length of the insertion. Lastly, *w*Bic contains a 104 bp insertion at bp position 647 that creates multiple downstream stop codons. These results suggest that the surface protein *wspB* has become psuedogenized at least four times in different *Wolbachia* lineages. Notably, the *Wolbachia* strains with a putatively pseudogenized version of *wspB* do not differ from the other strains in mean *Wolbachia* abundance in whole oocytes (Wilcoxon rank sum test, *W* = 36, *P* = 0.964), the posterior region (*W* = 37, *P* = 0.893), or the posterior cortex (*W* = 41, *P* = 0.622). This suggests that psuedogenization of *wspB* does not influence the diverse patterns of *Wolbachia* localization observed here. To gain insight into *wspB* cellular function, we ectopically expressed this gene (and the other *wsp* paralogs; WD_1063/*wsp* and WD_0489/*wspC)* in yeast using the Galactose expression system described above. Using this system, we found that ectopic expression of codon optimized *wspB* and *wsp* significantly inhibited yeast growth **(Supplemental Figure 9)**. Ectopic expression of *wspC* kills the cells. Based on similar ectopic expression studies (Sheehan et al., 2016), the suppression is likely due to *Wolbachia* proteins interacting with and disrupting the function of yeast components and cellular processes that are conserved and performing similar functions in insect cells.

Lastly, we ran similar phylogenetic analyses using the host phylogeny (as opposed to *Wolbachia*) to test whether host-associated factors might also contribute to the diverse *Wolbachia* distribution patterns in fly oocytes (**Supplemental Figure 10**). We generally found that *Wolbachia* localization patterns do not exhibit a phylogenetic signal on the host tree. *Wolbachia* abundance in whole oocytes (λ = 0.365 [0, 0.856], *P* = 0.656), at the posterior region (λ < 0.001 [0, 0.658], *P* = 1), and at the posterior cortex (λ < 0.001 [0, 0.637], *P* = 1) did not exhibit phylogenetic signal using the host phylogram, implying that host-associated factors are not as important in determining diverse *Wolbachia* localization patterns in oocytes.

### Vertical transmission in some host species likely relies on *Wolbachia* entering the germline from neighboring somatic tissues

Because *Wolbachia* strains with a Dispersed localization do not localize to the pole, one would expect maternal transmission to be compromised. However, previous studies demonstrate that some of these *Wolbachia*, such as the B-group strain *w*Mau in *D. mauritiana*, have high maternal transmission rates under standard laboratory conditions (i.e., constant 25°C), equivalent to that of *w*Mel in *D. melanogaster* (Meany et al., 2019; Hague et al., 2022). These findings suggest that, in addition to strict germline-to-germline transmission, *Wolbachia* may utilize alternative routes of maternal transmission. Because numerous vertically inherited endosymbionts are transmitted via cell-to-cell transmission from the soma to the germline (Frydman, 2006; Russell et al., 2019), we explored whether *Wolbachia* that infect *Drosophila* exploits this strategy.

To determine the developmental stage during which somatic *Wolbachia* might invade the germline in Dispersed species, we focused on B-group *w*Mau in *D. mauritiana.* Initially we examined the female germline (oocytes) of 3^rd^ instar larva for the presence of *Wolbachia*. For comparison, we also examined the oocytes of *w*Mel-infected *D. melanogaster* (a strain with a Posteriorly Localized concentration of *Wolbachia*) 3^rd^-instar larvae using anti-FtsZ to mark the *Wolbachia* and anti-Vasa to mark the germline. As shown in **Figure 5**, *Wolbachia* are abundantly present in ring-like patterns in many of the Vasa-positive cells in the center of the developing oocyte. Imaging the oocytes of *w*Mau-infected *D. mauritiana* revealed a similar pattern with *Wolbachia* concentrated in the developing female germline of 3^rd^ instar larva (**Figure 5**). Images of *w*Ri in *D. simulans* as well as *w*Bic in *D. bicornuta—*strains with a Dispersed localization and Posteriorly Clumped localization in mature oocytes, respectively—displayed the same ring-like structure in the developing oocyte of 3^rd^ instar larva, although they appeared to have a much higher titer than *w*Mau or *w*Mel (**Figure 5**). This indicates that in these species, in which vertical transmission occurs with a few or no *Wolbachia* present at the posterior site of germline formation in mature adult oocytes, *Wolbachia* are able to occupy the germline at some point between the final stages of oocyte maturation and development of the fertilized egg into a 3^rd^ instar larva. Presumably the source of the *Wolbachia* is via invasion from neighboring somatic cells.

**Figure 5.**
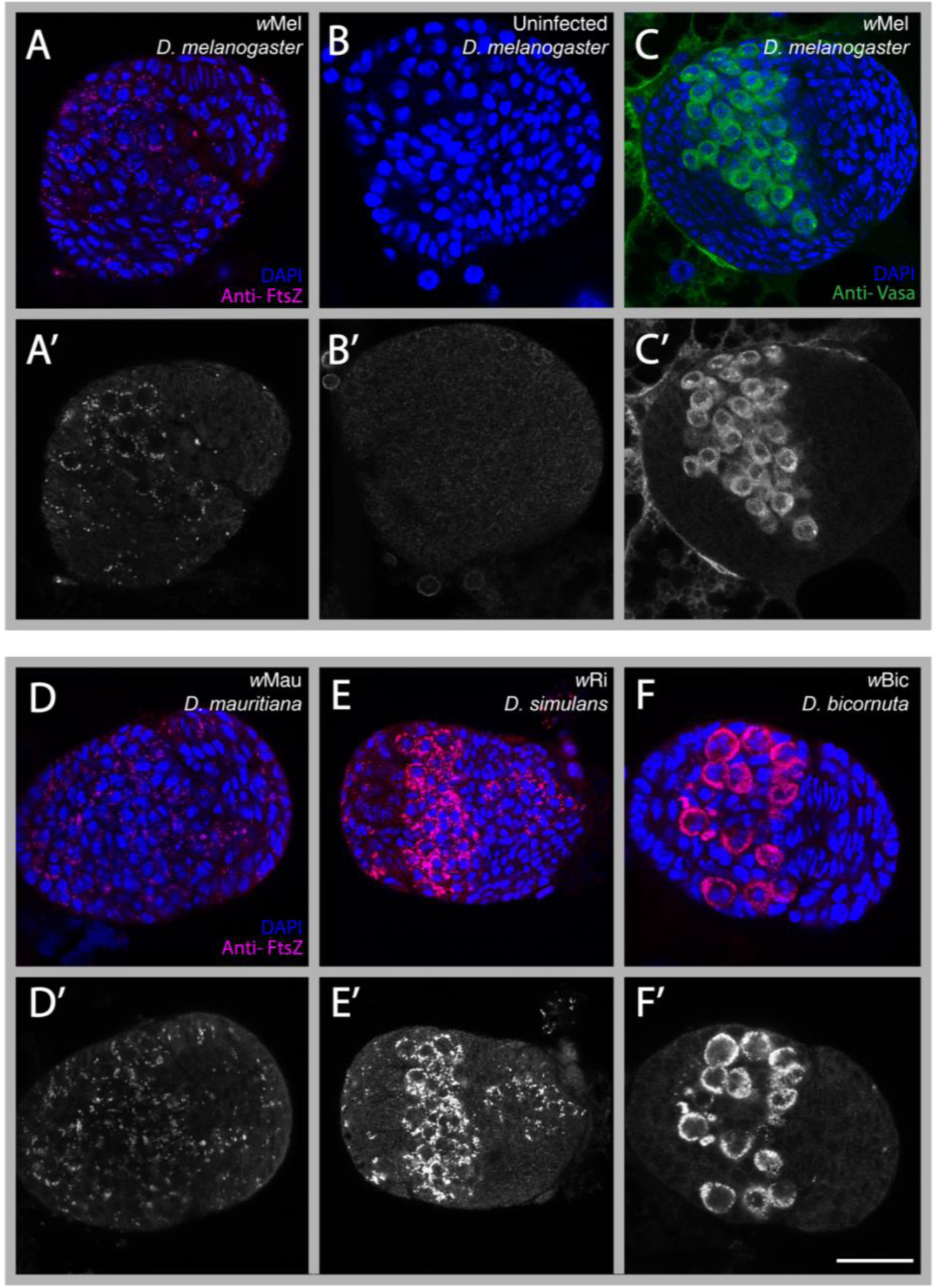
*Wolbachia*’s presence around the 3^rd^ instar larval germline, in multiple *Drosophila* hosts. *w*Mel *Wolbachia* stained with anti-FtsZ can be seen for *D. melanogaster*, in panel (A) shown in pink. Cells are marked with DAPI (blue). (A’) Grayscale channel of anti-FtsZ staining. (B) Uninfected ovary for *D. melanogaster*, (B’) grayscale channel for uninfected antibody binding. Germline location marked with anti-Vasa shown in green (C), grayscale channel (C’). *w*Mau *Wolbachia* localization in the ovary for *D. mauritiana* shown in panel (D), grayscale channel (D’). *w*Ri and *D. simulans* shown in panel (E), grayscale (E’). Panel (F), *w*Bic and *D. bicornuta*, (F’) grayscale channel. Scale bar shown at 25 uM.

To determine if *Wolbachia* occupied the germline prior to the 3^rd^ instar larval stage, we examined the newly formed germline of late blastoderm and cellularized embryos. At this stage, newly formed germline cells, known as the pole cells, form a distinct cluster of cells at the extreme posterior to the embryo. In all the four strains described above, using PI or anti-*Wolbachia* staining, *Wolbachia* are readily found in these posterior germline cells (**Figure 6**). Imaging *w*Mau in *D. mauritiana* early blastoderm embryos revealed the presence of *Wolbachia* in the posterior pole cells. This observation narrows the time window in which *Wolbachia* invades and occupies the germline from between the final stages of oocyte maturation to early blastoderm formation. Images taken of *D. mauritiana* in one-hour old embryos also showed *w*Mau positioned in the posterior pole, narrowing the developmental window of *Wolbachia* germline occupation between the final stages of oocyte maturation to one hour after fertilization **(Figure 6).** Interestingly, Veneti et al. (2004) reported that *w*Mau exhibits an anterior localization pattern during *D. mauritiana* embryogenesis; however, we found no such pattern for any of the embryos examined.

**Figure 6.**
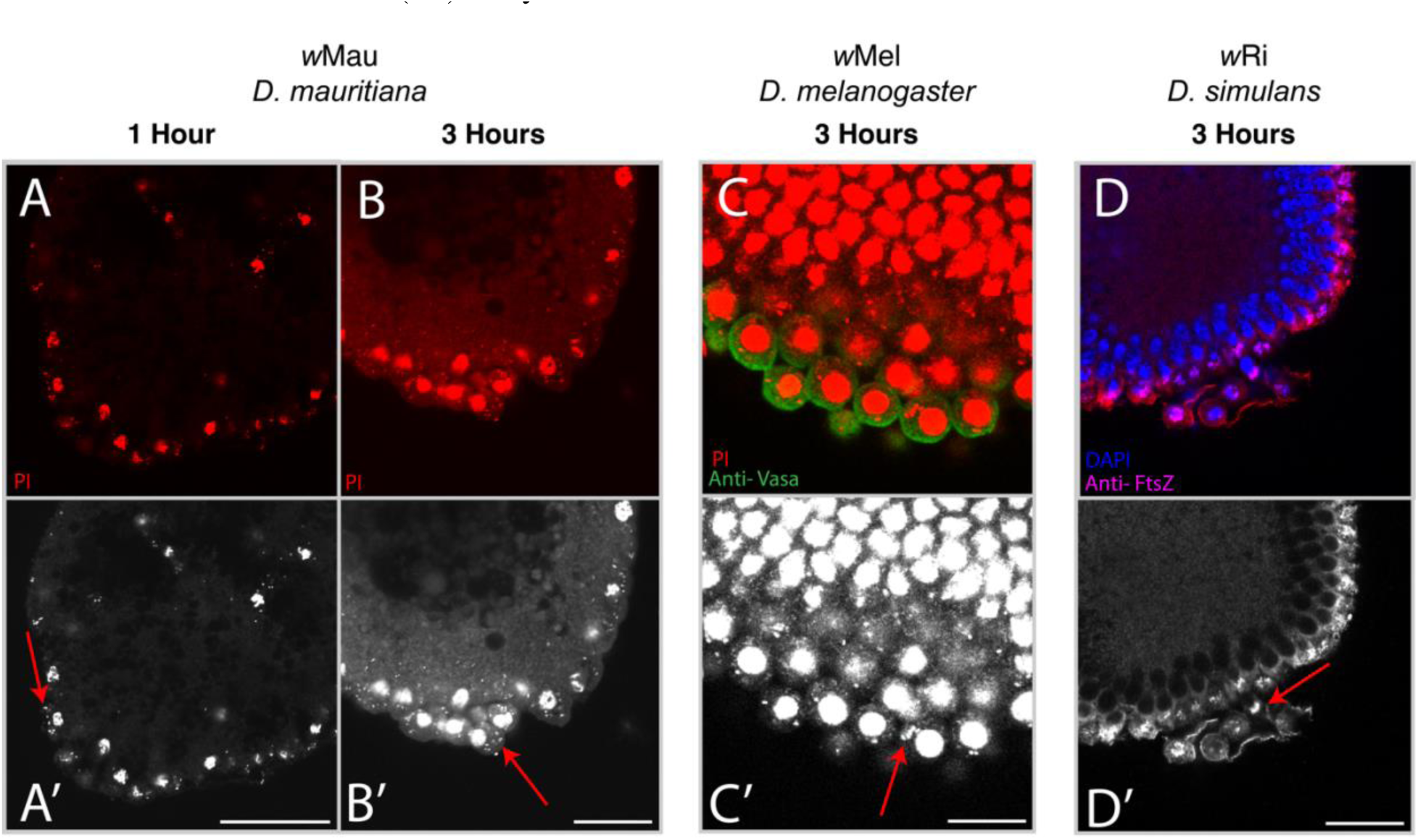
*Wolbachia*’s presence in the newly formed germline cells of late blastoderm and cellularized embryos. (A) 1 hour *D. mauritiana* (*w*Mau) embryo stained with PI. (A’) Greyscale of PI channel. (B) staining of 2-3 hour developed *D. mauritiana* (*w*Mau) embryo with PI. (B’) PI of greyscale channel. (C) PI and anti-Vasa (green) staining of *w*Mel *Wolbachia* in 2-3 hour *D. melanogaster* embryo. (C’) Greyscale of PI channel. (D) Anti-FtsZ/DAPI staining for *w*Ri *Wolbachia* in *D. simulans*. (D’) Greyscale of anti-FtsZ channel. Scale bars set at 25uM.

Our findings motivated us to examine *w*Mau in *D. mauritiana* oocytes just prior to egg deposition. Previous studies found that in many infected *Drosophila* species *Wolbachia* is concentrated in the most posterior positioned follicle cells (Kamath et al., 2018). These somatically-derived cells, known as polar cells, directly contact the oocyte posterior pole plasm. In accord with Kamath et al. (2018), we find that *Wolbachia* are sporadically distributed in subsets of more anterior positioned follicle cells encompassing the oocyte. *w*Mau presence in the polar follicle cells is a consistent feature in every *D. mauritiana* oocyte examined (**Figure 7**). Given their position, these *w*Mau-infected cells are likely the polar follicle cells. These observations raise the possibility that *Wolbachia* in the polar follicle cells may be a somatic source of germline *w*Mau in *D. mauritiana.* In spite of much effort, however, we have not been able to find cytological evidence for this hypothesis.

**Figure 7.**
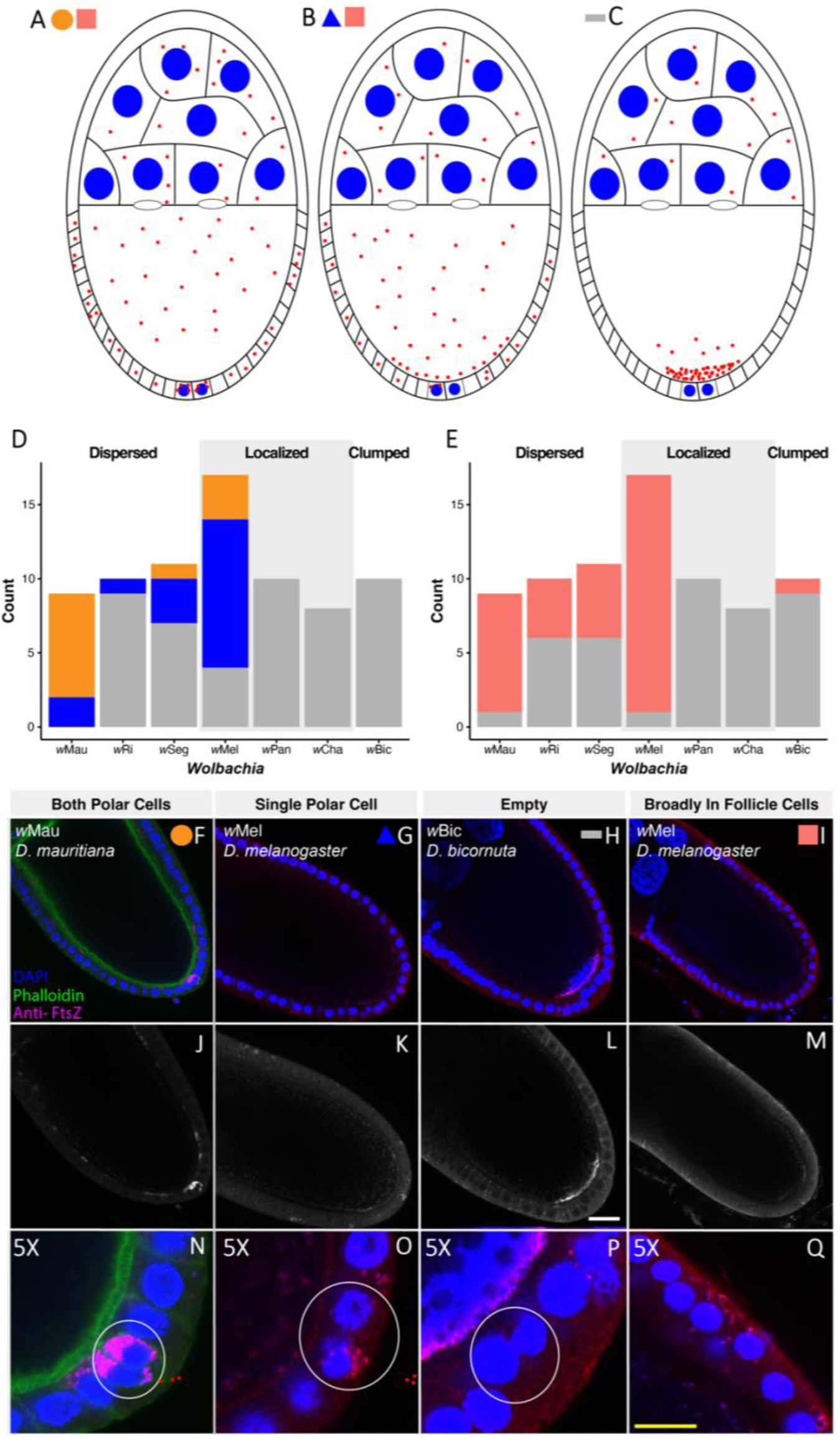
The A-group *w*Mel and B-group *w*Mau *Wolbachia* strains are frequently found in the polar follicle cells, as well as other follicle cells. (A-C) Schematic of categories used for description of follicle cell *Wolbachia*. (A) Dispersed localization pattern shown, both polar cells infected, with *Wolbachia* broadly present in follicle cells. (B) Posteriorly Localized pattern shown, one polar cell infected, with *Wolbachia* broadly present in follicle cells. (C) Posteriorly Clumped pattern shown, with no *Wolbachia* present in the polar cells or other follicle cells. (D) *Wolbachia* quantification in the polar follicle cells during oogenesis stages 9-10. *Wolbachia* were quantified in midsections of each oocyte that displays unique posterior localization patterns: Dispersed, Posteriorly Localized, and Posteriorly Clumped. For each strain, the count is shown for the number of oocytes observed with *Wolbachia* present in both polar follicle cells (orange), only one polar cell (blue), and neither polar cell (grey). (E) Counts of oocytes observed with *Wolbachia* present anywhere in the follicle cells (pink) or with *Wolbachia* absent (grey). (F) Representative anti-FtsZ staining of *w*Mau in *D. mauritiana* (Dispersed class) present in both polar follicle cells (corresponding to orange labels above). (G) Representative staining of *w*Mel in *D. melanogaster* (Posteriorly Localized class) present in only one polar cell (blue labels above). (H) Representative staining of *w*Bic in *D. bicornuta* (Posteriorly Clumped class) absent in both polar cells (grey labels above). (I) Representative staining of *w*Mel in *D. melanogaster* (Posteriorly Localized class) broadly present in anterior follicle cells (pink labeling above). (J-M) Greyscale anti-FtsZ channel of respective composite images above. (N-Q) 5X zoom of polar cells, with circles highlighting representative staining in the area. Scale bar set at 25uM.

To determine the extent to which *Wolbachia* polar follicle cell localization is conserved, we analyzed *Wolbachia* localization in the polar and more anterior follicle cells for seven *Wolbachia* strains with the following patterns: Dispersed (*N* = 3), Posteriorly Localized (*N* = 3), and Posteriorly Clumped (*N* = 1) (**Figure 7**). We scored for the presence of follicle cell *Wolbachia* in each oocyte (*N* = 75) using the following criteria: the presence of *Wolbachia* in either both or one of the posterior polar follicle cells (**Figure 7A**) and the presence of *Wolbachia* in the more anteriorly localized follicle cells (**Figure 7B**). This analysis revealed that in a high percentage of oocytes from Dispersed strains (a lack of the *Wolbachia* at the oocyte posterior), *Wolbachia* are frequently present in the posterior polar follicle cells. In contrast, with the exception of wMel, oocytes examined from strains in which *Wolbachia* are posteriorly concentrated (the Posteriorly Localized and Clumped strains), few possess *Wolbachia* in the posterior follicle cells. The seven *Wolbachia* strains exhibit significant variation in whether *Wolbachia* cells are found in the polar follicle cells (χ^2^_(12)_ = 65.15, *P* < 0.001) (**Figure 7A**). In particular, the divergent B-group strain *w*Mau stood out with a large proportion of oocytes containing *Wolbachia* in the two polar follicle cells. The A-group *w*Mel strain also had a large portion of oocytes with *Wolbachia* in one or two of the polar cells. We also found that the *Wolbachia* strains differ in whether *Wolbachia* cells are found in the other more anterior follicle cells (χ^2^_(6)_ = 43.29, *P* < 0.001) (**Figure 7B**). Again, the *w*Mau and *w*Mel strains had a large portion of oocytes with *Wolbachia* in the other more anterior follicle cells encompassing the oocyte. Of note, for *w*Mau in *D. mauritiana*, every oocyte had at least one of their polar cells filled with *Wolbachia*, observationally at a higher titer (**Figure 7**). This suggests that in the Dispersed strains, vertical transmission may be achieved via invasion of *Wolbachia* from the polar follicle cells, particularly for the divergent B-Group *w*Mau strain.

## DISCUSSION

Vertical transmission through the female lineage is a common strategy for many endosymbionts. These endosymbionts achieve high efficiencies of maternal transmission either through direct germline-to-germline transmission or invasion of the maternal germline through neighboring somatic cells (Russell et al., 2019). Because the former involves a continuous presence in the germline while the latter requires navigating a soma-to-germline passage, it is expected that each requires distinct interactions with and manipulations of host cellular processes. Thus, it would be expected that a given endosymbiont would have evolved to utilize one, but not both of these strategies. Here we explore this issue by examining cellular aspects of *Wolbachia* vertical transmission. In contrast to expectations, our comprehensive analysis demonstrates that *Wolbachia* likely rely on both strict germline-to-germline and soma-to-germline vertical transmission strategies. Which strategy employed varies among *Wolbachia* strains.

The initial anterior *Wolbachia* localization in the developing oocyte appears to be a conserved aspect of *Wolbachia* transmission, as all eight strains examined exhibit this localization. It is striking that this localization occurs prior to any known anterior determinant, indicating *Wolbachia* may associate with an as yet undiscovered host factor concentrated at the anterior. That this localization is conserved suggests that it has a functionally significant, but currently unknown role in *Wolbachia* transmission. While anteriorly localized, the amount of *Wolbachia* dramatically increases. Whether *Wolbachia* replication, transport from the nurse cells, or both are responsible for this increase is unknown. The anterior location may be a rich source of membrane for *Wolbachia* replication. The anterior localization also results in a high concentration of *Wolbachia* that closely associated with, and perhaps influences, the status of the oocyte nucleus.

To our surprise, we discovered that immediately after the release of *w*Mel *Wolbachia* from its anterior position in the oocyte (stage 6), the amount of *Wolbachia* in the oocyte dramatically decreases (see **Figure 1, Stage 6** and **Supplemental Figure 1**). This suggests that the *Wolbachia* are transported back into the nurse cell complex. The functional significance of this clearing of *Wolbachia* from the oocyte is unknown. The retreat of *Wolbachia* occurs during the time when the anteriorly positioned oocyte nucleus and neighboring follicle cells signal one another in order to establish the dorsal ventral axis (Merkle et al., 2020). Perhaps *Wolbachia* exits the oocyte in order to not disrupt this process. Given the orientation of microtubules at this stage, it is likely *Wolbachia* rapidly exits the oocyte through an association with the plus-end directed motor protein Kinesin.

Following microtubule reorientation and the return of *Wolbachia* to the oocyte, *Wolbachia* spreads posteriorly throughout the entire oocyte relying on the host motor protein Kinesin (Serbus and Sullivan, 2007). All of the *Wolbachia* strains examined are distributed toward the posterior regions suggesting the interaction with Kinesin is also conserved. How *Wolbachia* engages these motor proteins is unknown. Surprisingly, none of the known host Kinesin linker proteins are utilized by *Wolbachia,* suggesting *Wolbachia* may interact directly with Kinesin (Russell et al., 2018).

It is after this stage in which *Wolbachia* is distributed throughout the oocyte, that we observe variability among the *Wolbachia* strains, with each falling into one of three distinct classes: two distinct classes in which *Wolbachia* concentrate at the posterior (Posteriorly Localized and Posteriorly Clumped) and one class in which *Wolbachia* fail to exhibit a posterior concentration (Dispersed). The former two classes are distinguished by whether the vast majority (Posteriorly Clumped) or only a small fraction of the *Wolbachia* localize to the posterior pole (Posteriorly Localized). Here, our phylogenomic analyses revealed a strong correlation between *Wolbachia* posterior abundance and *Wolbachia* phylogenetic relationships. Generally, we find that closely related *w*Mel-like *Wolbachia* occur at higher abundance at the oocyte posterior, whereas *w*Ri-like *Wolbachia* exhibit a more dispersed distribution (**Figure 4**). These results suggest that factors intrinsic to *Wolbachia* help determine posterior localization patterns, which is consistent with previous *Wolbachia* transplantation studies demonstrating that posterior localization is determined by *Wolbachia* rather than host factors (Poinsot et al., 1998; Veneti et al., 2004; Serbus and Sullivan, 2007). Below, we discuss how specific factors might contribute to variation in *Wolbachia* localization patterns.

The Posteriorly Localized class, in which only a small fraction localize to the posterior pole, can be explained by previous work demonstrating that *w*Mel is a weak competitor for Kinesin; both *Wolbachia* and germplasm components rely on Kinesin for transport to the posterior pole (Russell et al., 2018). It is thought that *Wolbachia* has evolved to be a weak competitor because interference with the transport of essential germplasm components and thus germplasm formation may be disadvantageous for *Wolbachia* proliferation. The other posterior class (Posteriorly Clumped) is strikingly similar to that observed for *w*Mel in *D. melanogaster* in which the plus-end motor protein Kinesin is over-expressed and excess amounts of *Wolbachia* are transported to the posterior pole (Russell et al., 2018). It may be that the Posteriorly Clumped strains are much better competitors for host Kinesin. Alternatively, these strains could occur in host species with naturally much higher levels of host Kinesin, although our phylogenetic analysis did not support a role of host factors. Functional work will be required to formally test these hypotheses. In particular, it would be interesting to investigate whether there are instances in which *Wolbachia* disrupts the A-P axis (anterior-posterior) because of its accumulation at the posterior pole.

The posterior pole concentration among *Wolbachia* strains may also differ due to variability in the ability of *Wolbachia* to stably associate with the pole plasm. Previous studies demonstrated that *Wolbachia* posterior pole concentration requires intact pole plasm (Serbus and Sullivan, 2007). For example, in *oskar* mutations, a key pole plasm determinant, *w*Mel *Wolbachia* fails to accumulate at the posterior region (Hadfield and Axton, 1999; Serbus and Sullivan, 2007). As described, the oocyte experiences tremendous cytoplasmic streaming requiring posterior components to be anchored directly or indirectly to the cortex (Quinlan, 2016). This variability could be due to differences in the ability of *Wolbachia* to compete with other pole plasm components such as mitochondria for binding sites. Alternatively, the host species may vary in the extent to which their pole plasm accommodates *Wolbachia*. These alternatives can be explored through trans-infection studies.

It is likely that the composition of *Wolbachia* surface proteins play a major role in the extent of posterior localization. Our phylogenetic analyses revealed that the gene trees of two candidate surface proteins, particularly WD_1085, are strongly associated with *Wolbachia* localization patterns. The WD_1085 protein has sequence and structural homology to the bacterial outer membrane protein BamA, which consists of a large periplasmic domain attached to a 16-stranded β-barrel domain (**Supplemental Figure 11**). BamA is the central subunit of the β-barrel assembly machinery (BAM), which is essential for outer membrane protein biogenesis (Noinaj et al., 2013; Doyle and Bernstein 2019). Given this finding, it is intriguing that we find overexpression of BamA is sensitive to disruptions in microtubules, as these cytoskeletal elements are essential for outer membrane biogenesis in mitochondria (Mado et al., 2019). It may be that microtubules are also required for *Wolbachia* membrane biogenesis. While we highlight these loci as potential candidates, we note that our results should be interpreted with caution since the WD_1085 gene could be in linkage with other causal loci. Nonetheless, *Wolbachia* surface proteins are generally considered to be strong candidates for interactions with host cells (Werren et al., 2008; Baldridge et al., 2016; Hague et al., 2022).

Interestingly, our phylogenomic analyses did not find a correlation between host phylogenetic relationships and *Wolbachia* localization patterns, suggesting that host-associated factors are less important in determining *Wolbachia* posterior localization. This result contrasts with transplantation studies that indicate hosts may also play a significant role in determining *Wolbachia* abundance (Veneti et al., 2004). In addition, genome-wide screens reveal host factors play a major role in determining *Wolbachia* intracellular abundance (White et al., 2017; Grobler et al., 2018). Thus, variation in the host proteins that *Wolbachia* engages could still plausibly influence *Wolbachia* localization patterns. This would contrast with other endosymbionts such as *Listeria*, which relies on a surface protein that binds and polymerizes host actin to propel the bacteria within and between host cells (Kühn and Enninga, 2020). Because the interaction between ActA and actin is essential for cell-to-cell transmission, natural variants of this interaction and transmission strategy have not been discovered. In addition to *Wolbachia* and host factors, nutrients and other environmental factors have also been shown to influence *Wolbachia* abundance and localization in host oocytes for closely related *w*Mel-like *Wolbachia* strains (Serbus et al., 2015; Hague et al., 2020a; Hague et al., 2022).

Two classes in which *Wolbachia* concentrates in the posterior pole (Posteriorly Localized and Posteriorly Clumped) provide a ready explanation for the cellular mechanisms by which it is vertically transmitted through generations. In every generation, *Wolbachia* targets the site of germline formation in the developing oocyte. This also explains why strains exhibiting this pattern, such as *w*Mel, exhibit efficient transmission strategies under typical laboratory conditions (Hague et al., 2022). The third, Distributed class, that exhibits no-to-low *Wolbachia* in the posterior germplasm of the mature oocyte is puzzling. This would be expected to result in embryos lacking *Wolbachia* in the germline. However, within hours after fertilization, *Wolbachia* is clearly present in the germlines of both sexes (**Figure 6**). This implies that either these *Wolbachia* strains are capable of invading the germline from neighboring somatic cells or a few undetected *Wolbachia* are delivered late in oogenesis to the posterior pole. Support for the first hypothesis comes from the fact that in filarial nematodes *Wolbachia* germline invasion from the soma has been directly observed (Landmann et al., 2012). While germline invasion via cell-to-cell transmission has not been directly observed in insects, a number of lines of evidence indicate that it likely occurs. In *Drosophila* cell culture, *Wolbachia* efficiently undergoes cell-to-cell transmission (White et al., 2017; White et al., 2017). *Wolbachia* injected into the abdomen of adult *Drosophila* females migrate to and occupy the germline and follicle stem cells (Frydman et al., 2006). Collections directly from nature reveal some strains with developing oocytes lacking *Wolbachia* (Casper-Lindley et al., 2011). In these strains, all of the mature oocytes were infected, suggesting an alternate route of infection via neighboring somatic cells.

As previously hypothesized, given the posterior follicle cells are directly adjacent to the pole plasm, we propose *Wolbachia* present in these follicle cells may be the source of germline *Wolbachia* (Kamath et al., 2018). Our results are in accord with and complement previous studies demonstrating *Wolbachia* concentrated in the posterior polar follicle cells in a number of *Wolbachia*-infected species (Kamath et al., 2018). For example, in *w*Mau-infected *D. mauritiana*, a Dispersed class, there is a striking concentration of *Wolbachia* in the posterior follicle pole cells directly adjacent to the pole plasm. Thus, *Wolbachia* in these somatic cells are well positioned to invade the pole plasm becoming incorporated into the germline of the next generation (**Figures 7 and 8**). In contrast, in the Posteriorly Localized and Clumped strains, we found no to few *Wolbachia* in the posterior polar follicle cells. We note that invasion of *Wolbachia* from posterior follicle cells into the germline would require *Wolbachia* to cross the seemingly impenetrable newly formed vitelline membrane, which could suggest alternative sources of germline *Wolbachia* (we thank the reviewer for this insight). Nonetheless, taken together the data strongly argue for a somatic-to-germline route of vertical transmission in some *Drosophila* species.

**Figure 8.**
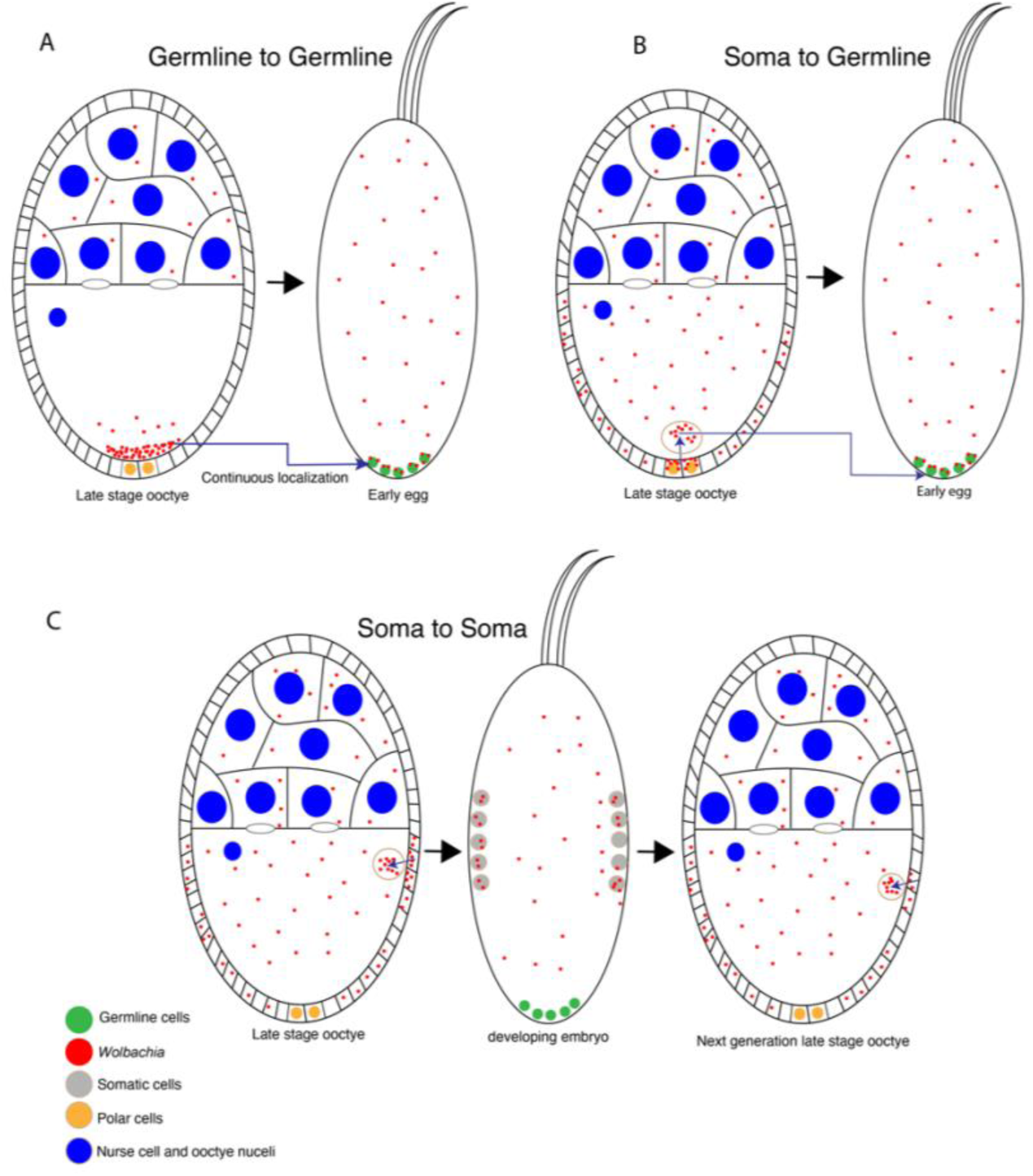
Multiple routes of *Wolbachia* vertical transmission. (A) In *Wolbachia* strains that concentrate at the posterior pole plasm, vertical transmission likely occurs through a continuous maintenance in the germline. (B) In strains where no or few *Wolbachia* localize to the posterior pole plasm, vertical transmission likely occurs through *Wolbachia* invading the germline from neighboring somatic cells**. (**C) In theory, vertical transmission could occur through a symbiont invading the oocyte from neighboring somatic cells never associating with the germline.

Among the *Wolbachia* strains examined, *w*Mel stands out as an exception. Not only is it concentrated at the posterior pole of the oocyte, but there is also a large *Wolbachia* concentration in the posterior polar follicle cells. Thus, *w*Mel may maintain both robust germline-to-germline and soma-to-germline modes of transmission. Accordingly, it has a high efficiency of vertical transmission under laboratory conditions (but see Hague et al. 2022). Notably, we found that *w*Mel and *w*Mau are also both quite prevalent in the more anteriorly located follicle cells of stage 10 oocytes (**Figure 7**), raising the possibility that vertical transmission could even occur without *Wolbachia* ever directly associating with the host germline (i.e., soma-to-soma transmission; **Figure 8**).

Surprisingly, literature surveys reveal that germline invasion of endosymbionts from the soma every generation is the most common form of vertical inheritance (Russell et al., 2019). Vertical transmission via a continuous presence in the germline is much less common. One explanation is that hosts maintains mechanisms preventing endosymbiont occupation during the formative stages of germline development. In instances in which the endosymbiont is maintained in the germline, this may evolve into an obligate relationship in which development of the germline depends on the presence of the endosymbiont (Sullivan, 2017). Examples of this include the leafhopper (*E. plebeus*), in which the endosymbiont is required for normal embryonic development (Sander, 1968). In addition, removal of *Wolbachia* from some insects results in increased apoptosis and abnormal oocyte development (Dedeine et al., 2001; Pannebakker et al., 2007). Here we find that *Wolbachia* exhibit evidence of both the soma-to-germline and germline-to-germline transmission strategies (**Figure 8**), suggesting that *Wolbachia* may stand out as a rarity among endosymbionts.

## MATERIALS AND METHODS

### Drosophila Stocks

All stocks were grown on standard brown food (Sullivan et al., 2000) at 25°C with a 12h light/dark cycle. Uninfected *Drosophila* stocks were generated by tetracycline-curing of the infected stock (Serbus et al., 2015). Listed in Table 1 are the 19 different *Wolbachia* strains we examined, which infect 18 different *Drosophila* host species. This included six *w*Mel-like strains (Cooper et al., 2019; Hague et al., 2020a), five *w*Ri-like strains (Turelli et al., 2018), seven other A-group *Wolbachia*, and the B-group strain *w*Mau that diverged from A-group *Wolbachia* up to 46 million years ago (Meany et al., 2019). Each genotype was generated as an iso-female line by sampling a single gravid female from the field and placing her individually into a vial.

### Ovary, embryo and larva fixation and staining

Newly eclosed flies were transferred to fresh food for 3-5 days for aging. Approximately 10 females were dissected for each slide preparation. Ovaries were fixed using a modification of previously published procedures (Brendza et al., 2000; Russell et al., 2018). Ovaries were removed and separated in phosphate buffer solution and then fixed in a 200 μl devitellinizing solution (2% paraformaldehyde and 0.5% v/v NP40 in 1x PBS) mixed with 600 μl heptane for 20 min at room temperature, on a shaker. After removal of the organic layer by brief centrifugation for sample isolation, the ovaries were washed 3 times with PBS-T (0.1% Triton X-100 in 1x PBS) along with three five-minute washes. Samples were treated with RNAse A (10 mg/ml) and left overnight at room temperature. Samples were then washed again a few times with PBS-T and then stained with a dilute solution of phalloidin for actin staining on a rotator for 1 hour. Samples were washed again with 3 quick PBS-T rounds and then solution changes with PBS-T over two hours. 60uL of propidium iodide (PI) in mounting media was added to the sample after removal of the wash solution and left again overnight. Ovaries were then mounted and carefully separated out again for ease of imaging and removal of excess mature eggs. Slides were coated in nail polish and stored at -20℃ until imaged.

Embryos were collected for 1-3 hours on plates and fixed in equal volume 32% paraformaldehyde and heptane and fixed as previously described (Sullivan et al., 2000). After extraction with methanol, embryos were blocked for one hour in 5% PBST-BSA (2.5g bovine serum albumin fraction V in 50ml PBS-T) at room temperature. Embryos were then incubated with anti-WD_0009 or anti-Ftfz at 1:500 overnight at 4℃. After three washes with 1% PBST-BSA over one hour, embryos were incubated in Alexa 488 Anti-Rabbit at 1:500 for one hour at room temperature. Samples were then washed three more times over one hour with 1% PBST-BSA. Embryos were mounted with media containing DAPI and stored at -20℃ until imaged.

Larval fat bodies were dissected and fixed from 3^rd^ instar larvae using a modified version of a published protocol (Maimon and Gilboa, 2011). Fat bodies were dissected in 1X PBS then fixed in 5% paraformaldehyde in PBS for 20 minutes with gentle agitation. Fat bodies were washed for five minutes, 10 minutes, then 45 minutes with cold 1% NP40 in PBS. After washes, fat bodies were blocked in 5% PBST-BSA (2.5g bovine serum albumin fraction V in 50ml PBS-T) for one hour at room temperature. Fat bodies were then incubated with anti-FtsZ at 1:500 overnight on a rocker at 4℃, washed three times over one hour with 1% PBST-BSA, then incubated with Alexa 488 Anti-Rabbit at 1:500 at room temperature for one hour. After three additional washes over one hour with 1% PBST-BSA, fat bodies were mounted onto a slide with media containing DAPI. Larval ovaries were isolated from fat bodies using 0.1mm needles before sealing the slide with nail polish. Samples were then stored at -20℃.

### Confocal microscopy

Imaging was performed on an inverted Leica DMI6000 SP5 scanning confocal microscope. Optical sections of a Nyquist value of 0.38 were used. A variety of zooms were used to optimize image viewing, with most being set at 1.5X. Propidium iodide was excited with the 514 and 543 nm lasers, and emission from 550 to 680 nm was collected. GFP was imaged with the 488 nm laser, and emission from 488 to 540 nm was collected. Alexa 633 was imaged with the 633 laser, and emission from 606 to 700 nm was collected. All imaging was performed at room temperature. Images were acquired with Leica Application Suite Advanced Fluorescence software.

### *Wolbachia* quantification and analysis

Following Russell et al. (2018), we used the polygon selection tool to select three different regions of the oocyte (see Fig. S1 in Russell et al. 2018): the whole oocyte, the posterior region, and the posterior cortex. The “area” and “integrated density” were measured for each region. Additionally, the average of a quadruplicate measure of “mean gray value” beside the stained oocytes was measured as background fluorescence. The <u>C</u>orrected <u>T</u>otal <u>C</u>ell <u>F</u>luorescence (CTCF) was calculated as integrated density – (area x background mean gray value) for each region.

All statistical analyses were performed in R. We first tested whether our three qualitative classes (Posteriorly Clumped, Posteriorly Localized, Dispersed) differed significantly in quantitative estimates of *Wolbachia* fluorescence (CTCF). We used a one-way ANOVA to test if the three groups differed based on their mean CTCF values from the whole oocyte, the posterior region, and the posterior cortex. Distribution and leverage analyses indicated that a transformation of *x*’ = log(*x* +1) was needed to meet assumptions of normality. We tested for significance using an *F* test with type III sum of squares using the “Anova” function in the *car* package (Fox and Weisberg, 2019). We conducted Tukey multiple comparisons among the three groups using the “TukeyHSD” function.

### Genomic data

In order to perform the phylogenomic analysis and generate phylograms, we obtained *Wolbachia* sequences from publicly available genome assemblies, which included *w*Mel (Wu et al., 2004), *w*Ri (Klasson et al., 2009), *w*Ha (Ellegaard et al., 2013), *w*Aura, *w*Tria, *w*Pan (Turelli et al., 2018), *w*Ana (Salzberg et al., 2005), and *w*Rec (Metcalf et al., 2014). All other *Wolbachia* sequences included in this study (*w*Bai, *w*Bic, *w*Boc, *w*Burl, *w*Cha, *w*Curt [*w*Tsa], *w*Nik, *w*Seg, *w*Tris, *w*Leo) were obtained using Illumina sequencing and previously described methods (Hague et al., 2020b). Briefly, tissue samples for genomic data were extracted using a DNeasy Blood & Tissue kit (Qiagen). DNA was cleaned using Agencourt AMPure XP beads (Beckman Coulter, Inc.) following manufacturers’ instructions, and eluted in 50 μl 1 × TE buffer for shearing. DNA was sheared using an E220 Focused Ultrasonicator (Covaris Inc.) to a target size of 400 bp. We prepared libraries using NEBNext® Ultra™ II DNA Library Prep with Sample Purification Beads (New England BioLabs). We indexed samples using NEBNext® Multiplex Oligos for Illumina® (Index Primers Set 3 & Index Primers Set 4), and 10 μL of each sample were shipped to Novogene (Sacramento, CA, USA) for sequencing using Illumina HiSeq 4000, generating paired-end 150 bp reads.

We obtained publicly available host sequences for *D. melanogaster* (Hoskins et al., 2015), *D. simulans* (Hu et al., 2013), *D. ananassae* (Clark et al., 2007), *D. pandora* (Turelli et al., 2018), *D. mauritiana* (Meany et al., 2019), *D. auraria, D. triaurari*, *D. baimaii*, *D. bicornuta* (aff. *bicornuta*), *D. bocki*, *D. burlai*, *D. chauvacae* (cf. *chauvacae*), *D. curta* (aff. *tsacasi*), *D. nikananu*, *D. seguyi*, *D. tristis*, and *D. leontia* (Conner, 2021). A *D. recens* genome assembly was kindly provided by Kelly Dyer and Rob Unckless. We note that three of the *montium* host species listed above have had recent updates to their taxonomic nomenclature: *D. bicornuta* has been renamed to aff. *bicornuta*, *chauvacae* has been renamed to cf. *chauvacae*, and *curta* has been renamed to aff. *tsacasi*. See Conner et al. (2021) for a full discussion of *montium* species names and relationships.

### *Wolbachia* phylogenetic analysis

Raw Illumina reads from the newly sequenced *Wolbachia* strains were trimmed using Sickle version 1.33 (Joshi and Fass, 2011) and assembled using AbySS version 2.0.2 (Jackman et al., 2017). *K* values of 71, 81, and 91 were used, and scaffolds with the best nucleotide BLAST matches to known *Wolbachia* sequences with *E*-values less than 10^-10^ were extracted as the draft *Wolbachia* assemblies. For each strain, we chose the assembly with the highest N50 and the fewest scaffolds (**Supplemental Table 2**). The newly assembled genomes and the previously published assemblies were annotated using Prokka version 1.11, which identifies homologs to known bacterial genes (Seemann, 2014). To avoid pseudogenes and paralogs, we only used genes present in a single copy with no alignment gaps in all of the genome sequences. Genes were identified as single copy if they uniquely matched a bacterial reference gene identified by Prokka. By requiring all homologs to have identical length in all of the *Wolbachia* genomes, we removed all loci with indels. A total of 66 genes totaling 43,275 bp met these criteria. We then estimated a Bayesian phylogram using RevBayes 1.0.8 under the GTR + Γ + I model partitioned by codon position (Höhna et al., 2016). Four independent runs were performed, which all converged on the same topology. All nodes were supported with Bayesian posterior probabilities of 1.

Similar methods were used to generate gene trees for candidate *Wolbachia* loci putatively involved in host interactions during oogenesis. This included the three *wsp* paralogs (WD_1063 [*wsp*/RS04815], WD_0009 [*wspB*/RS00060], WD_0489 [*wspC*/RS06475]) and three other *Wolbachia* surface proteins (WD_1085 [bamA/RS04910], WD_0501 [p44/MSP2/RS0225], WD_1041 [peptidase M2/RS04710]). Here, the WD_XXXX locus tags refer to the *w*Mel genome assembly (Wu et al., 2004).The protein-coding sequences for each locus were extracted from the assemblies using BLAST and the *w*Mel reference sequences. Notably, we found evidence of insertions in the WD_0009 (*wspB*) sequences of *w*Boc, *w*Leo, and *w*Bic (see Results). We removed these large insertions from the alignment to generate the *wspB* gene tree. In addition, BLAST identified WD_0489 (*wspC*) in *w*Mel-like *Wolbachia* and closely related *w*Bai, *w*Leo, and *w*Boc, but yielded no hits in the other *Wolbachia* strains (*w*Nik, *w*Tsa, *w*Pan, *w*Ana, *w*Tria, *w*Aura, *w*Ri, *w*Mau), indicating that *wspC* is either too diverged from the *w*Mel reference for BLAST recognition or alternatively *wspC* is only present in *w*Mel-related *Wolbachia* strains. Sequences for each gene were aligned with MAFFT 7 (Katoh and Standley, 2013). We then used RevBayes and the GTR + Γ + I model partitioned by codon position to generate gene trees for each locus.

### Host phylogenetic analysis

Host phylogenies were generated using the same nuclear genes implemented in Turelli et al. (2018): aconitase, aldolase, bicoid, ebony, enolase, esc, g6pdh, glyp, glys, ninaE, pepck, pgi, pgm, pic, ptc, tpi, transaldolase, white, wingless, and yellow. We used BLAST with the D. melanogaster coding sequences to extract orthologs from the genomes of each host species. Sequences were then aligned with MAFFT 7. Finally, we used RevBayes and the GTR + Γ + I model partitioned by codon position and gene to accommodate potential variation in the substitution process among genes, as described in Turelli et al. (2018). Our initial phylogenetic analysis produced a polytomy in the melanogaster species group regarding the relationships among the montium, melanogaster, and ananassae subgroups (**Supplemental Figure 10**). These relationships have been resolved in previous genomic studies involving additional taxa (Turelli et al., 2018; Suvorov et al., 2022); therefore, we generated a constrained host tree in RevBayes that enforced previously defined relationships among the three subgroups: ((montium, melanogaster), ananassae).

### Tests for phylogenetic signal

The resulting phylograms were used to test whether cellular *Wolbachia* abundance in oocytes exhibits phylogenetic signal on either the *Wolbachia* or host phylogenies. We used our quantitative estimates of *Wolbachia* abundance to test for phylogenetic signal using Pagel’s lambda (λ) (Pagel, 1999). Here, we used total *Wolbachia* fluorescence in whole oocytes, the posterior region, and the posterior cortex (log-transformed CTCF) as continuous characters to calculate maximum likelihood values of Pagel’s λ. A Pagel’s λ of 0 indicates that character evolution occurs independently of phylogenetic relationships, whereas λ = 1 is consistent with a Brownian motion model of character evolution. We used the “fitContinuous” function in GEIGER (Harmon et al., 2008) and a likelihood ratio test to compare our fitted value to a model assuming no phylogenetic signal (λ = 0). We also used a Monte Carlo-based method to generate 95% confidence intervals surrounding our estimates using 1,000 bootstrap replicates in the *pmc* package (Boettiger et al., 2012). When applicable, we conducted additional analyses to evaluate whether the number of taxa in our phylogeny (*N* = 19 *Wolbachia* strains, *N* = 18 host species) limited our ability to detect significant departures from λ = 0 (Hague et al., 2020b; Hague et al., 2021). Small phylogenies are likely to generate near-zero values simply by chance, not necessarily because the phylogeny is unimportant for trait evolution (Boettiger et al., 2012). To evaluate whether larger phylogenies increase the accuracy of estimation, we simulated trees with an increasing number of *Wolbachia* strains/host species (*N* = 25, 50) and our empirical estimates using the “sim.bdtree” and “sim.char” functions in the *geiger* R package (Harmon et al., 2008). We then re-estimated confidence intervals using the larger simulated trees.

Our tests for phylogenetic signal using the constrained host phylogram are presented in the main text; however, we note that our analysis using the unconstrained host tree (with a polytomy at the base of the *melanogaster* species group) produced similar results. Using the unconstrained tree, we found that *Wolbachia* abundance in whole oocytes (λ = 0.782 [0, 0.987], *P* = 0.469), the posterior region (λ = 0.951 [0, 1], *P* = 0.377), and the posterior cortex (λ < 0.001 [0, 0.563], *P* = 1) did not exhibit significant phylogenetic signal.

### Structural analysis of *Wolbachia* candidate genes

We used HHPred to identify protein domains of WD_1085 and WD_0501 using the COPe70_2.07, Pfam-A_v33.1, COG_KOG_v1.0, and SMART_v6 databases (Zimmermann et al., 2018; Gabler et al., 2020). Here we used default setting, including an E-value cutoff for MSA generation of 1e-3, a minimum coverage of MSA hits of 20%, and minimum probability in hitlist of 20%, and a maximum of 250 target hits. As expected, both WD_1085 and WD_0501 yielded multiple hits of varying length for domains associated with outer membrane proteins that form beta barrel structures (results summarized in **Supplemental Table 3**). We used SignalP-6.0 to identify the signal peptide cleavage sites for WD_1085 between aa residues 25 and 26 (Teufel et al., 2022) and removed the signal peptide for homology modeling. SignalP was unable confidently identify the signal peptide sequence for WD_0501, so we used the full sequence for homology modeling. We then used I-TASSER to model the three-dimensional structure of each protein through sequence homology (Yang and Zhang 2015; Zhang et al., 2017). For WD_1085, we used PDB:4K3B with an amino acid sequence identity of 23% in the threaded aligned region as a template (Noinaj et al., 2103). For WD_0501, we used PDB:2MLH with an amino acid sequence identity of 17% in the threaded aligned region as a template (Fox et al., 2014). PyMOL was used for image generation (Schrodinger 2015). Results are displayed in **Supplemental Figure S11**.

### Ectopic expression of *Wolbachia* surface proteins in yeast

SC-Leucine yeast media was prepared as described previously (Rose et al., 1990). Where present, benomyl was added to media from a 10mg/ml stock in DMSO to a final concentration of 2.5 ug/ml. The yeast expression assays are a modified version of the protocol described by Sheehan et al. (2016). *Wolbachia* genes were codon optimized for expression in *S. cerevisiae* and synthesized by Twist Bioscience (South San Francisco). The *wsp*, *wspB* and *wspC* genes were gifts of the Free Genes project (https://stanford.freegenes.org/). These genes were inserted into pGH448 using the New England Biolabs Golden Gate kit according to the manufacturer’s instruction. This placed the *Wolbachia* genes under control of the *GAL1* promoter and *CYC1* transcriptional terminator. This plasmid, which will be described in more detail elsewhere, carries a CEN/ARS element and also contains the kanamycin resistance and *LEU2* genes for selection in bacteria and yeast respectively.

The indicated plasmids were transformed into yeast strain GHY1934 [genotype] and GHY1551 [genotype]. Both of these strains are isogenic to S288C and are *GAL2+*. Transformed cells were grown to saturation in SC-Leucine liquid media containing 2% glucose. These cells were counted, diluted to 1×10^-7^ cells per ml, five-fold serial diluted and spotted to SC-Leucine plates with either 2% glucose or 2% galactose as the carbon source and incubated at the times and temperatures indicated in the figure legends.

## DATA AVAILABILITY STATEMENT

Newly generated genomic data are available on GenBank (BioProject PRJNA907328). All other data are publicly available on Dryad (doi:10.5061/dryad.v6wwpzh0q).

## ACKNOWLEDGEMENTS

We thank Dr. Benjamin Abrams (UCSC Life Sciences Microscopy Center, RRID: SCR_021135) for his technical support and assistance with microscopy experiments. We thank Dr. Shelbi Russell for their helpful advice on imaging protocols and analysis. We thank Dr. Irene Newton for kindly supplying the anti-FtsZ antibody. We thank Will Conner for guidance with bioinformatic analyses and Tim Wheeler for laboratory assistance. We thank Drew Endy and the Free Genes project for the gift of *Wolbachia* genes and advice on construction of pGH448. Research reported in this publication was supported by the National Institute of General Medical Sciences of the National Institutes of Health under Award Number R35GM139595 to W. Sullivan and R35GM124701 to B.S. Cooper. This work was also supported by the National Science Foundation (NSF) grant (1456535) to W. Sullivan and an NSF CAREER Award (2145195) to B.S. Cooper. We thank the Big Creek Reserve for use of their facilities. The microscopic analysis was enabled through the generous support of the IBSC microscopy facility at UC Santa Cruz.

## SUPPLEMENTAL INFORMATION

**Supplemental Figure 1.**
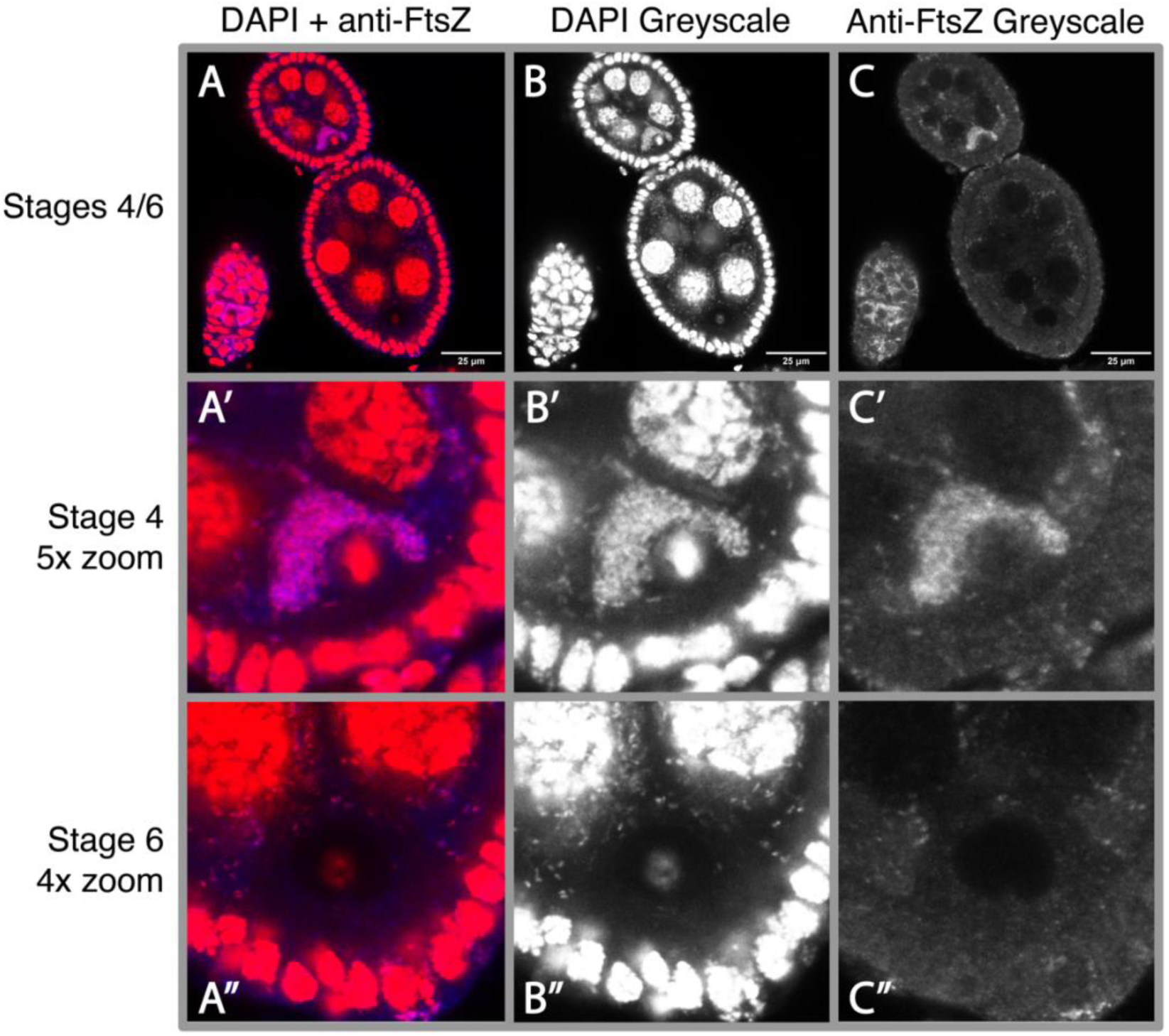
(A) Representative image of the reduction of *w*Mel *Wolbachia* titer between stages 4/5 and 6 in *D. melanogaster* oogenesis. *Wolbachia* stained with anti-FtsZ shown in pink. Scale bars set at 25 uM. (B) Greyscale DAPI channel. (C) Greyscale of anti-FtsZ channel. (A’-C’) 5X zoom around the nucleus of the oocyte during stage 4 shown for each channel. (A’’-C”) 4X zoom around the nucleus during stage 6 for each respective channel seen above. At least 10 oocytes across 3 slides were imaged with Z-stacks for a visual description of this trend.

**Supplemental Figure 2.**
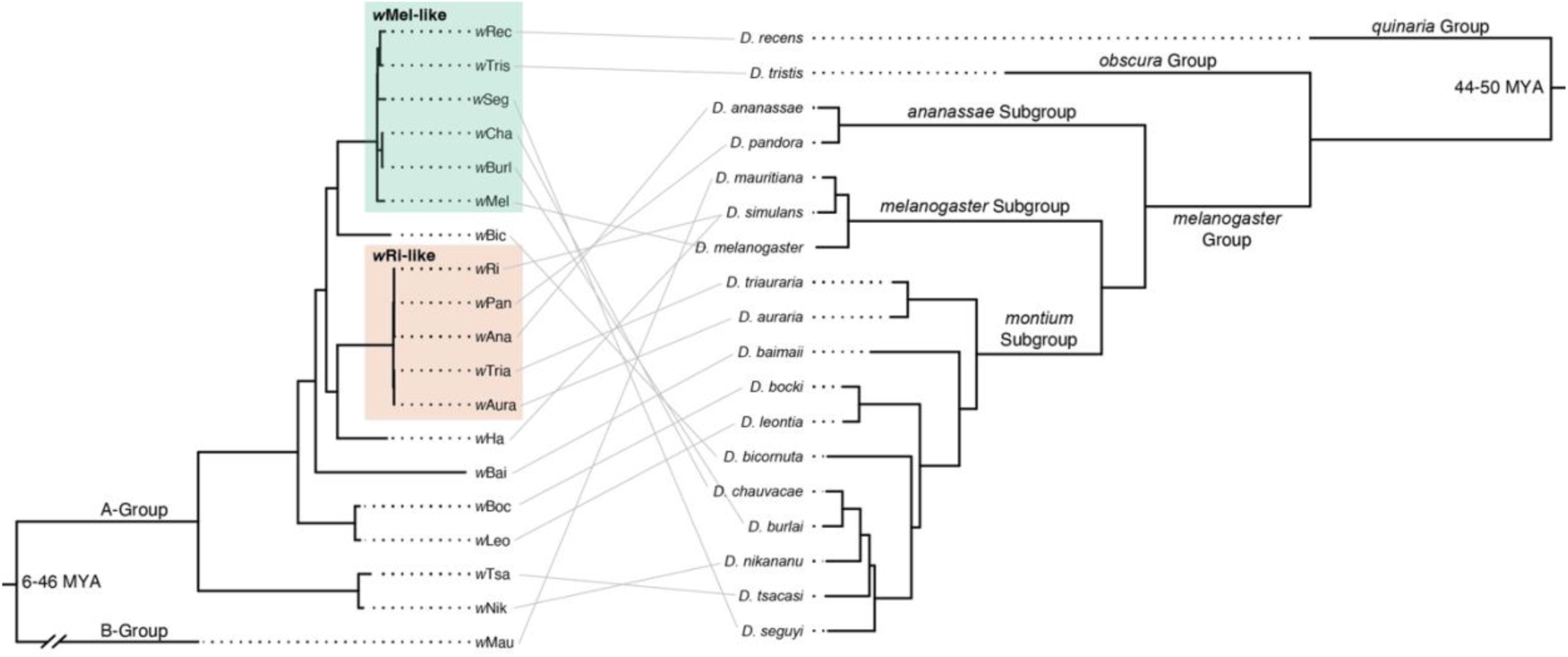
Bayesian phylograms depicting the evolutionary relationships among *Wolbachia* strains diverged up to 46 million years (left) and *Drosophila* host species diverged up to 50 million years (right). *w*Mel- and *w*Ri-like clades of closely related *Wolbachia* are labeled on the left. Grey lines pair *Wolbachia* strains with their *Drosophila* host species, which highlights patterns of topological discordance due to introgressive and horizontal transfer of *Wolbachia* among host species. The *Wolbachia* phylogram was estimated using 66 full-length and single-copy genes (43,275 bp) of equal length. All nodes have posterior probabilities >0.95. The host phylogram was estimated using 20 conserved single-copy genes (see Materials and Methods regarding host tree topology). *Wolbachia* and host divergence times in millions of years (MYA) are reproduced from Meany et al. (2019) and Suvorov et al. (2022), respectively.

**Supplemental Figure 3.**
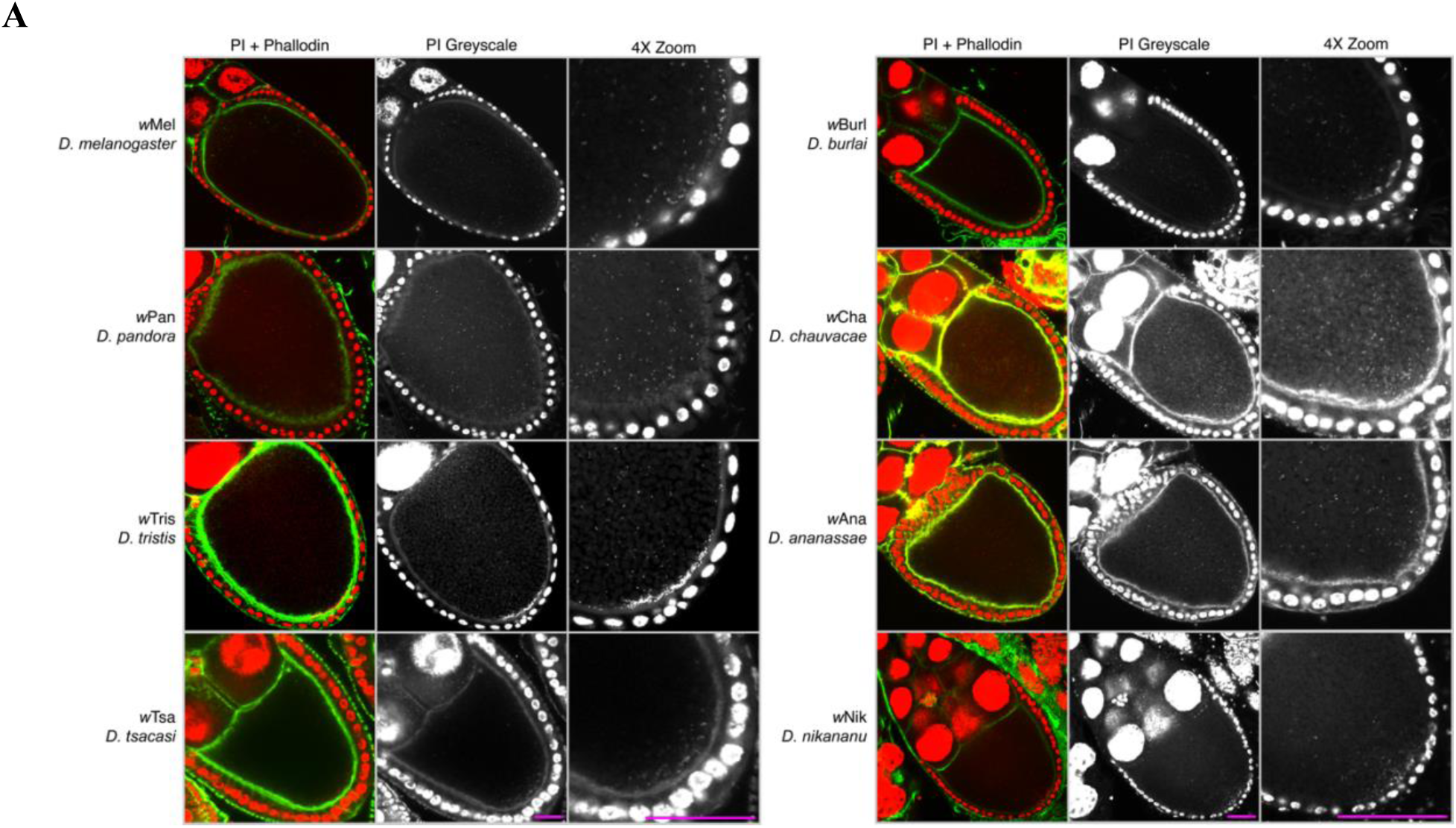
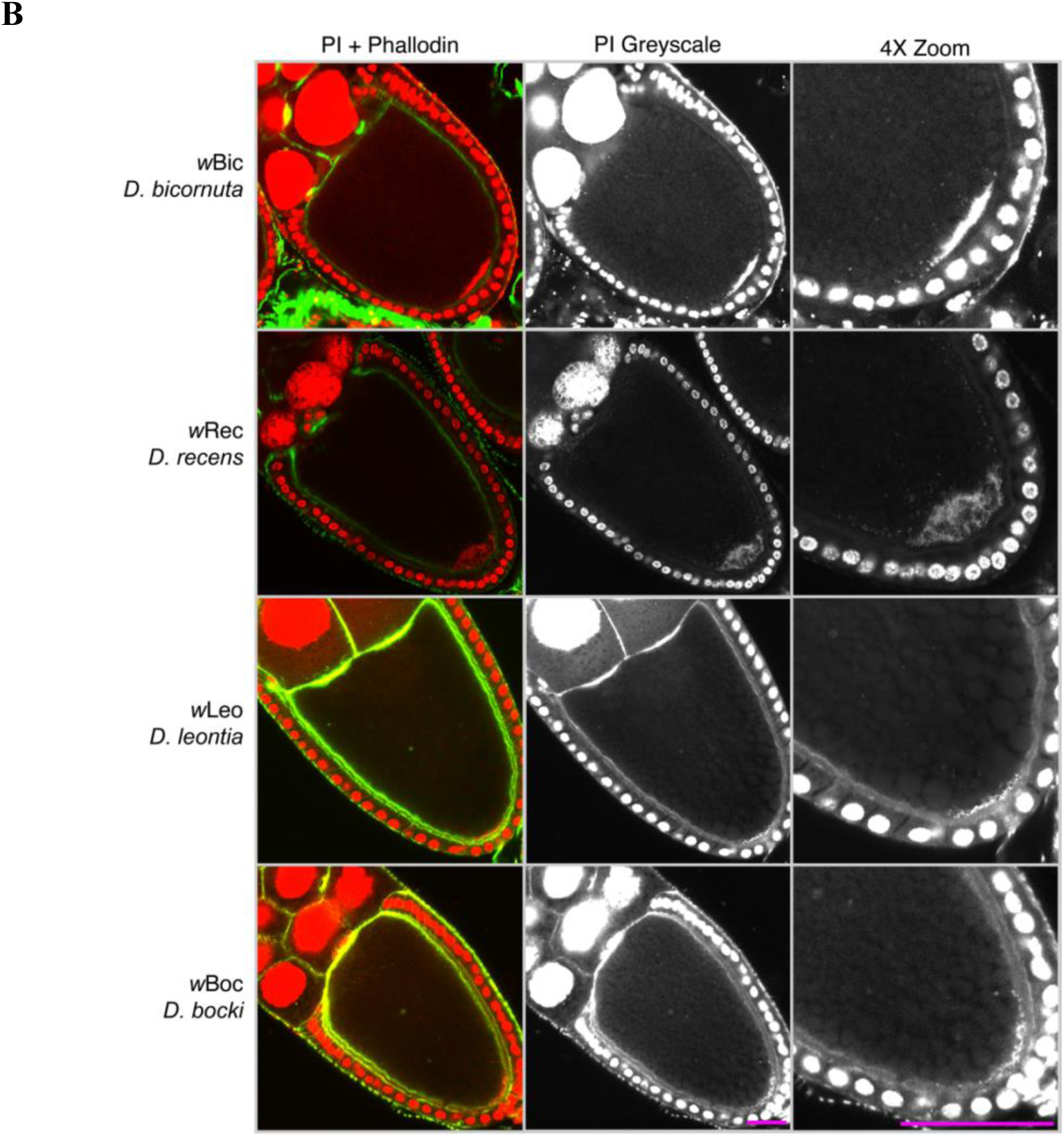
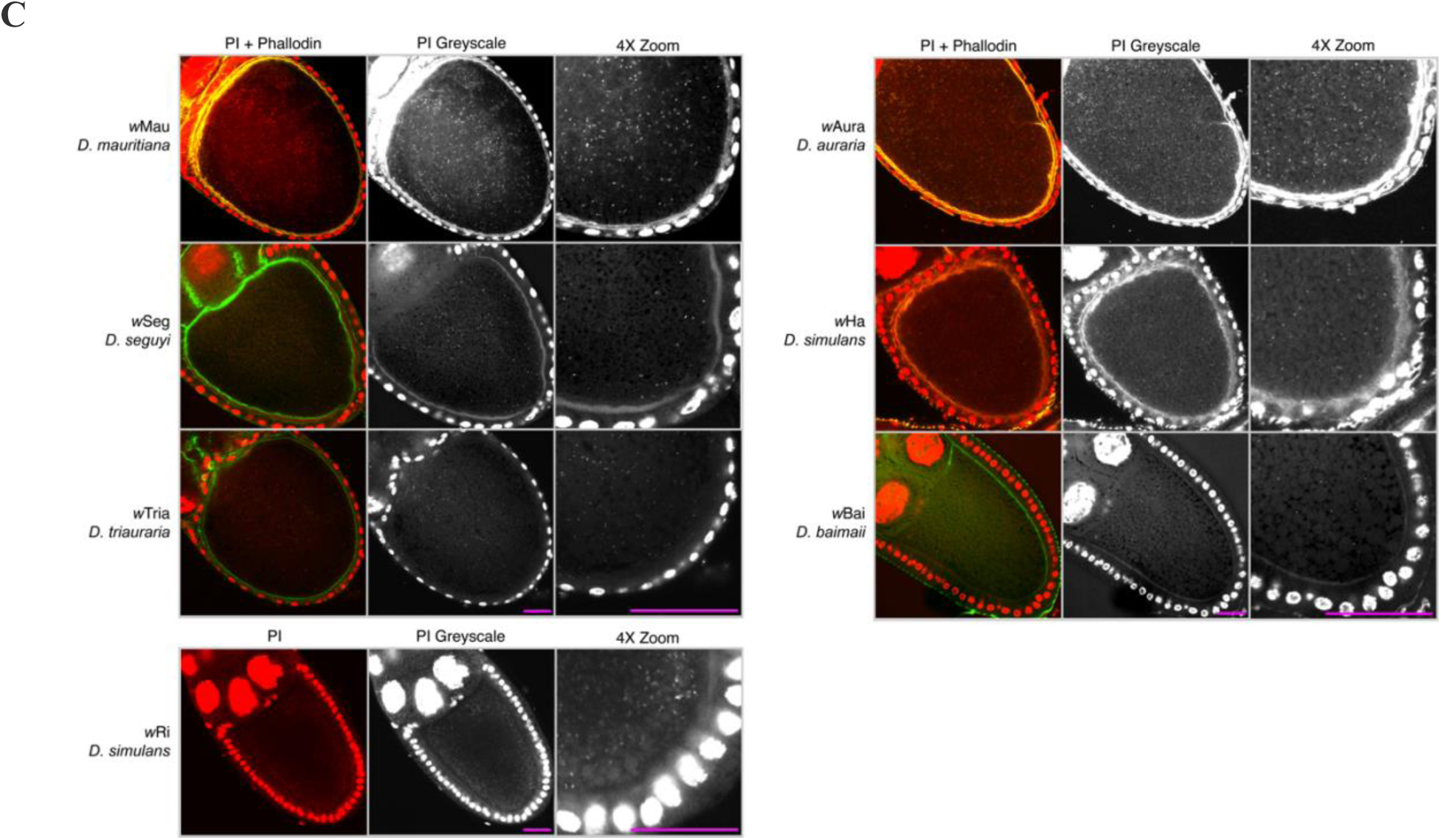
Representative images of *Wolbachia* posterior localization patterns. Confocal micrographs of *Drosophila* oocytes DNA-stained with PI (red) and actin-stained with phalloidin (green) show representative examples of infected *Drosophila* species with different *Wolbachia* localization classes. Second columns depict single greyscale channel images of PI staining. Third columns depict an enlarged PI-stained image of the posterior region of each oocyte. Panels A, B and C are grouped into *Wolbachia* strains that exhibit a Posteriorly Localized, Posteriorly Clumped, or Dispersed localization pattern respectively. Scale bar set at 25 uM.

**Supplemental Figure 4.**
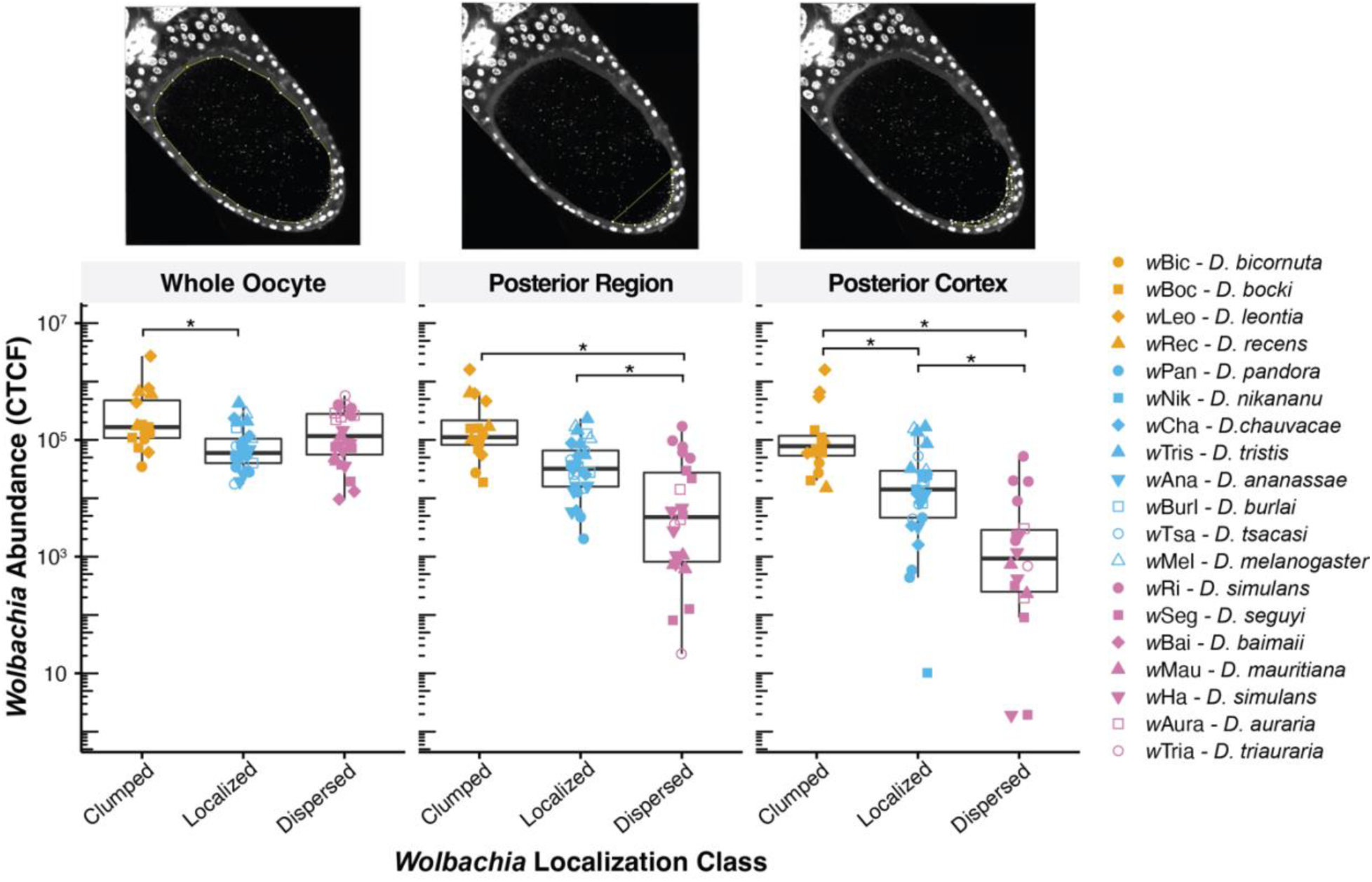
Cellular *Wolbachia* abundance in stage 10 oocytes measured as *Wolbachia* fluorescence due to propidium iodide (CTCF). Points are color coded by *Wolbachia* localization class (Posteriorly Clumped, Posteriorly Localized, Dispersed), with unique shapes indicating each *Wolbachia* strain and host species. Asterisks indicate significant differences among groups based on one-way ANOVAs and Tukey comparisons using *P*-values <0.05 adjusted for multiple comparisons. Above, a schematic is shown for fluorescent quantification of different regions of the oocyte using the PI greyscale channel: the whole oocyte, the posterior region, and the posterior cortex.

**Supplemental Figure 5.**
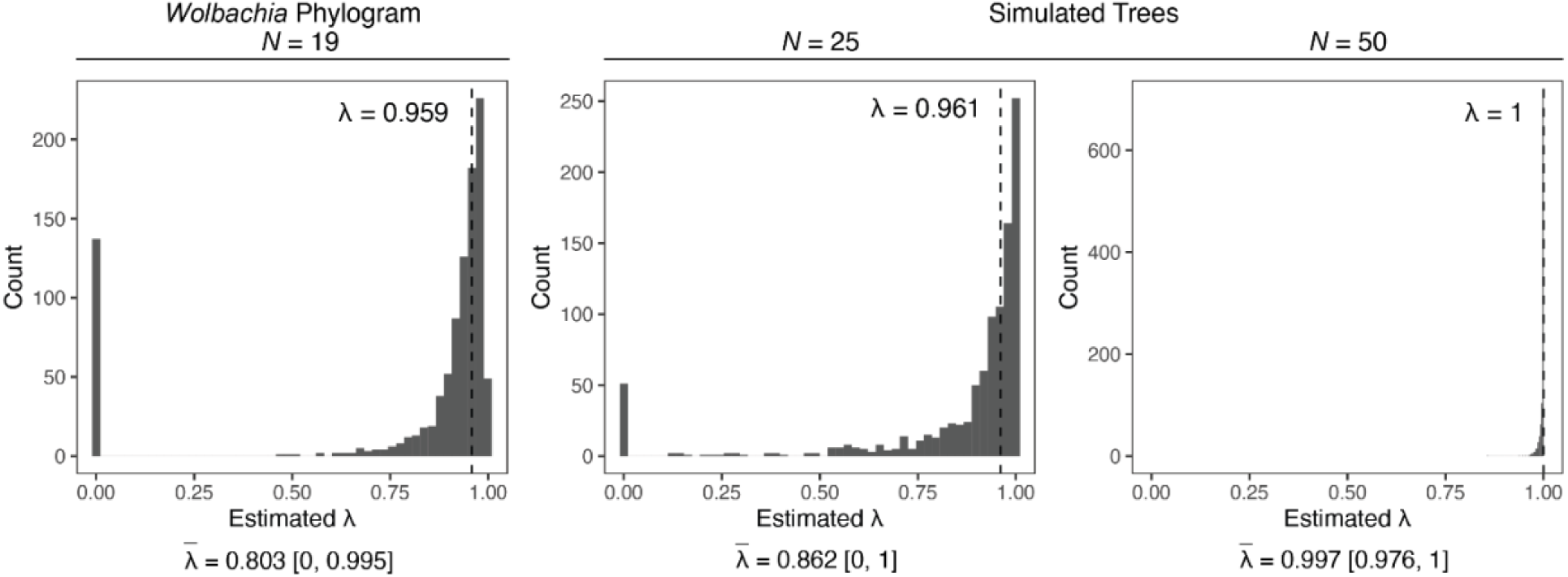
Distribution of maximum likelihood estimates of *λ* from 1,000 bootstrap replicates based on the *Wolbachia* phylogram and *Wolbachia* abundance at the oocyte posterior cortex (log-transformed CTCF). The bootstrap analysis of our *Wolbachia* phylogram is shown to the left (*N* = 19 *Wolbachia* strains). To the right are simulated phylogenies with an increasing number of *Wolbachia* strains included (*N* = 25, 50). For simulated trees, character evolution was simulated with our *λ* estimate of 0.959 using the “sim.bdtree” and “sim.char” functions in the *Geiger* R package (Harmon et al. 2008). For each graph, fitted *λ* values for the original phylogeny are shown above with a vertical dashed line. Note that fitted *λ* values for the simulated phylogenies differ slightly from *λ* = 0.959, because “sim.char” uses a Brownian-motion model to simulate character evolution along the phylogeny. Below each graph, the mean estimate of *λ* from the 1,000 replicates 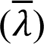 is shown with associated 95% confidence intervals. Small phylogenies (e.g., *N* = 19) are likely to generate many near-zero *λ* values by chance, not necessarily because the phylogeny is unimportant for trait evolution (Boettiger et al. 2012). As the number of strains in our analysis increases (*N* = 25, 50), bootstrapped estimates of *λ* cluster around the true *λ* value fitted to the original phylogeny.

**Supplemental Figure 6.**
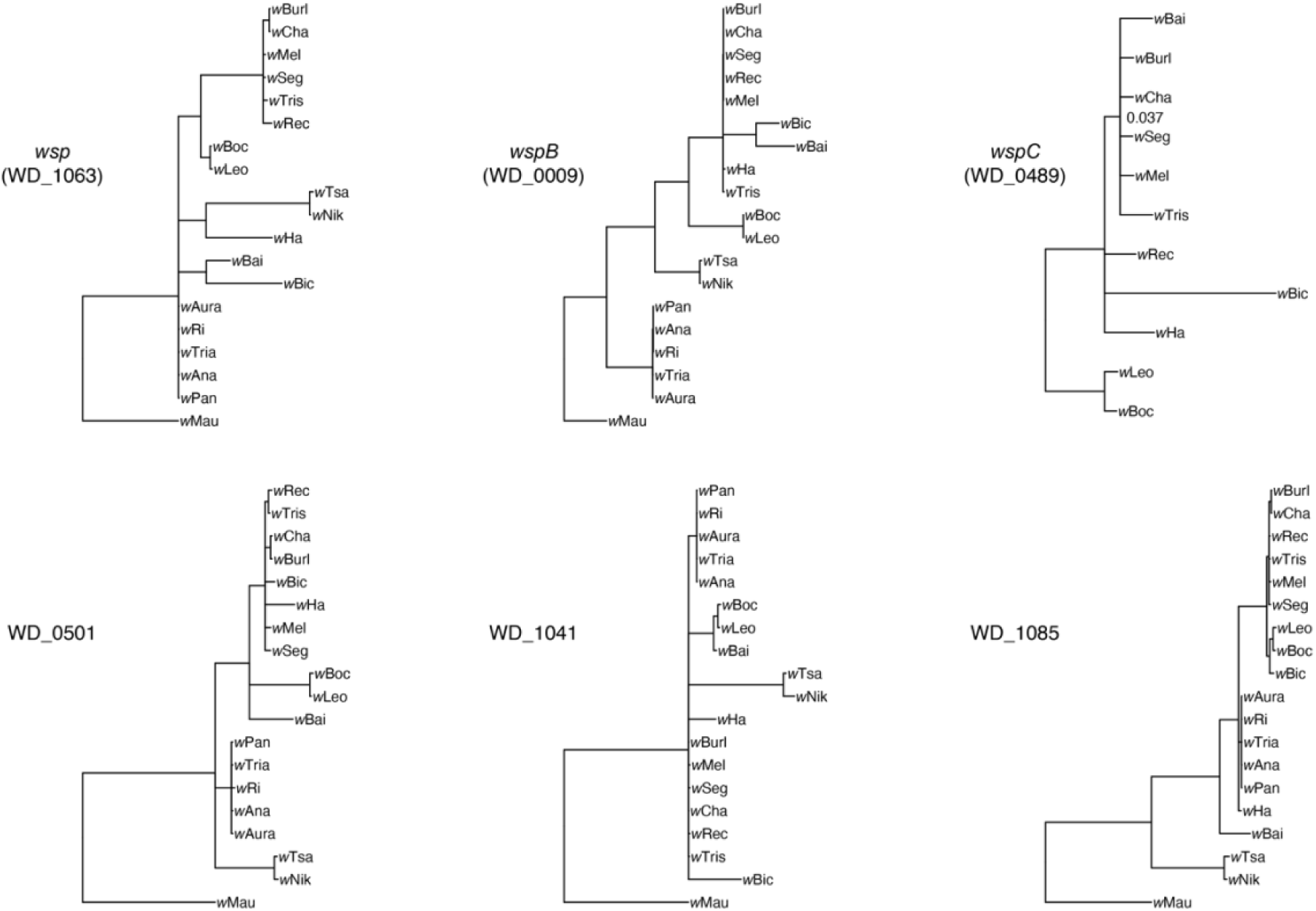
Gene trees of *Wolbachia* surface proteins. All nodes have Bayesian posterior probabilities of >0.95 unless otherwise noted (see only *wspC*).

**Supplemental Figure 7.**
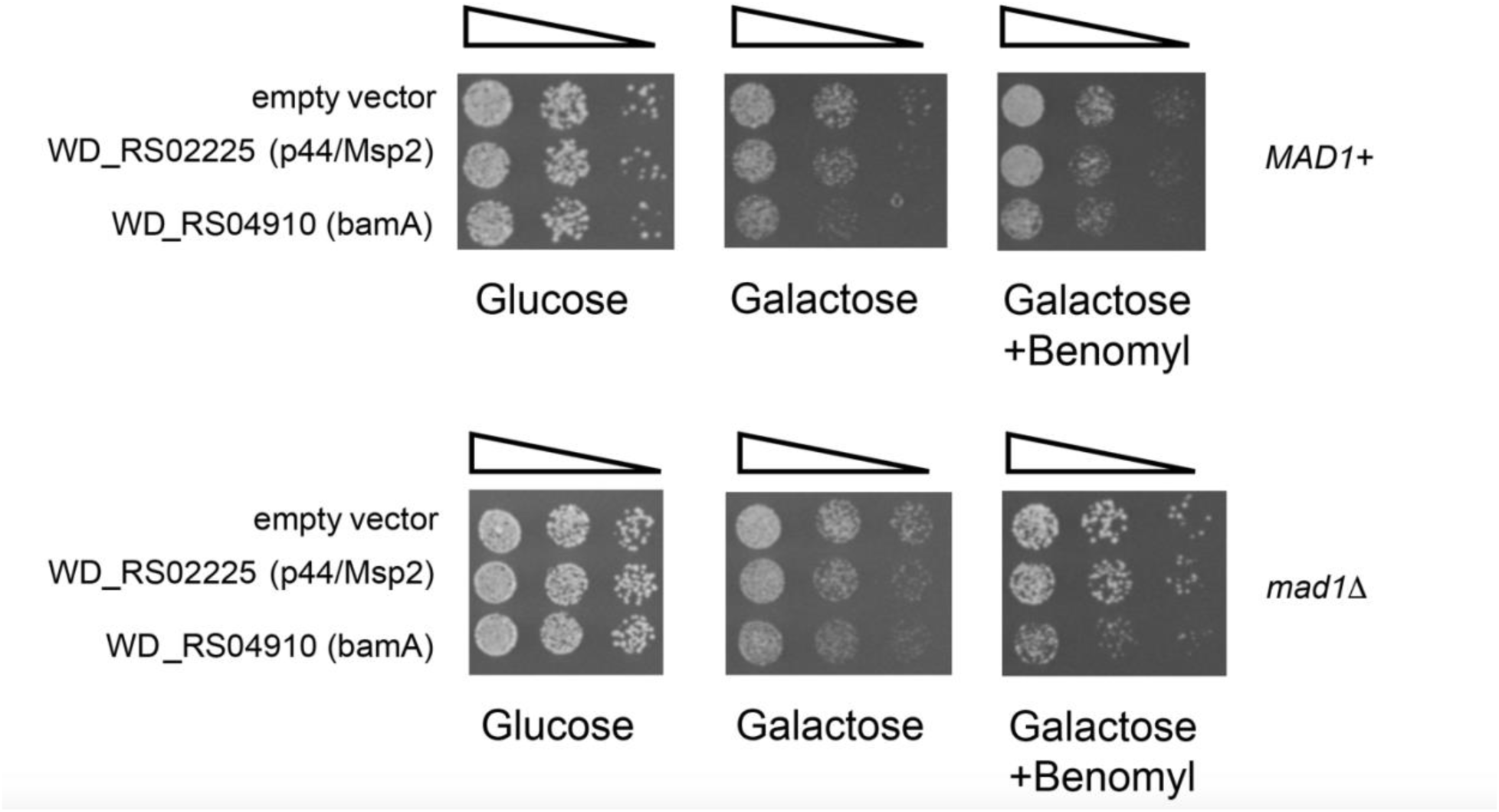
To gain insight into WD_0501 (RS0225/p44/Msp2) and WD_1085 (RS04910/bamA) cellular function, we ectopically expressed these genes in yeast using a Galactose inducible promoter. Under normal growth conditions, ectopic expression of both genes inhibited growth, with WD_1085 (RS04910/bamA) exhibiting a more pronounced inhibition. Growth inhibition of WD_1085 was dramatically increased when the integrity of microtubules was compromised with the yeast placed in the mad1 spindle assembly checkpoint background or the microtubules were compromised directly through the addition of benomyl. These results suggest that WD_1085 may directly or indirectly interact with host microtubules.

**Supplemental Figure 8.**
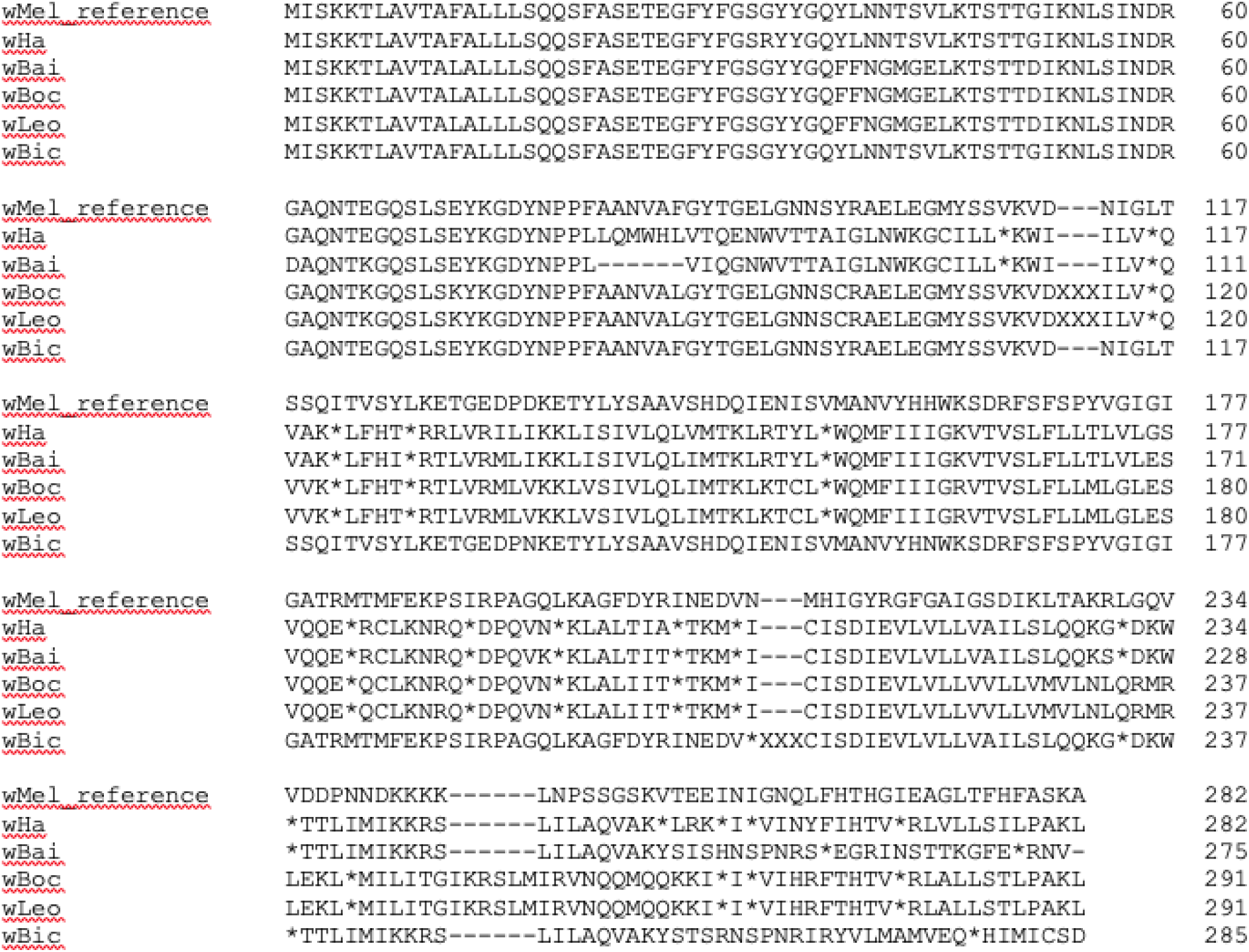
Full amino acid alignment of *wspB* (WD_0009) homologs in *w*Mel and the *Wolbachia* strains with a putatively pseudogenized version of the surface protein. Stop codons are indicated by asterisks and insertions of varying length are indicated by “XXX”.

**Supplemental Figure 9.**
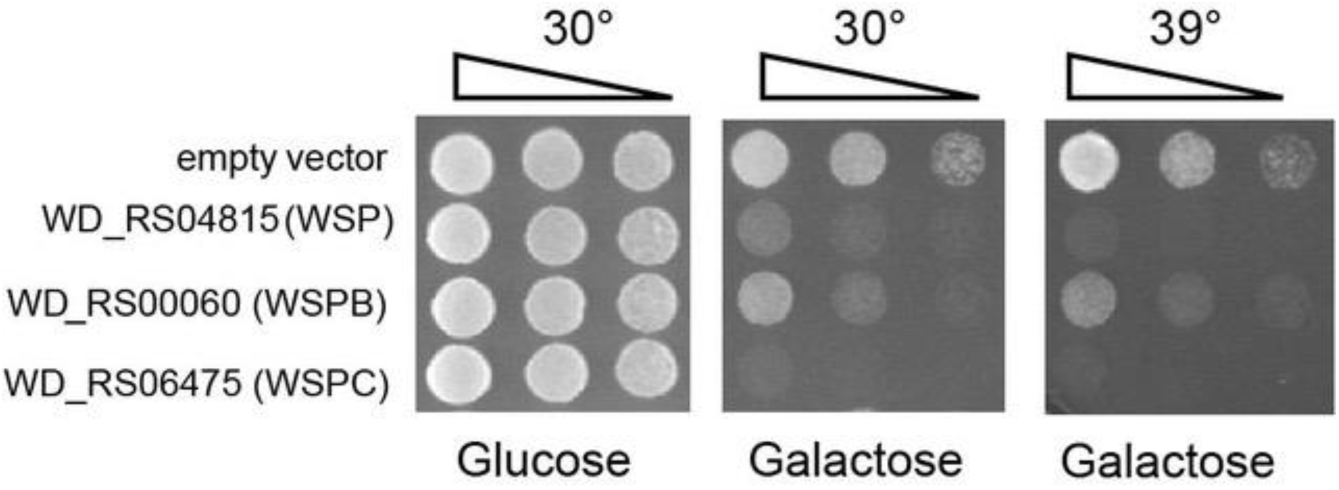
Predicted *Wolbachia* surface proteins WD_1063 (*wsp*/RS04815), WD_0009 (*wspB*/RS00060), and WD_0489 (*wspC*/RS06475) were assayed for their effect on eukaryotic cellular processes through ectopic expression in yeast. Note that galactose induction of gene expression of *wspC* kills the cells, while expression of *wsp* and *wspB* inhibits growth.

**Supplemental Figure 10.**
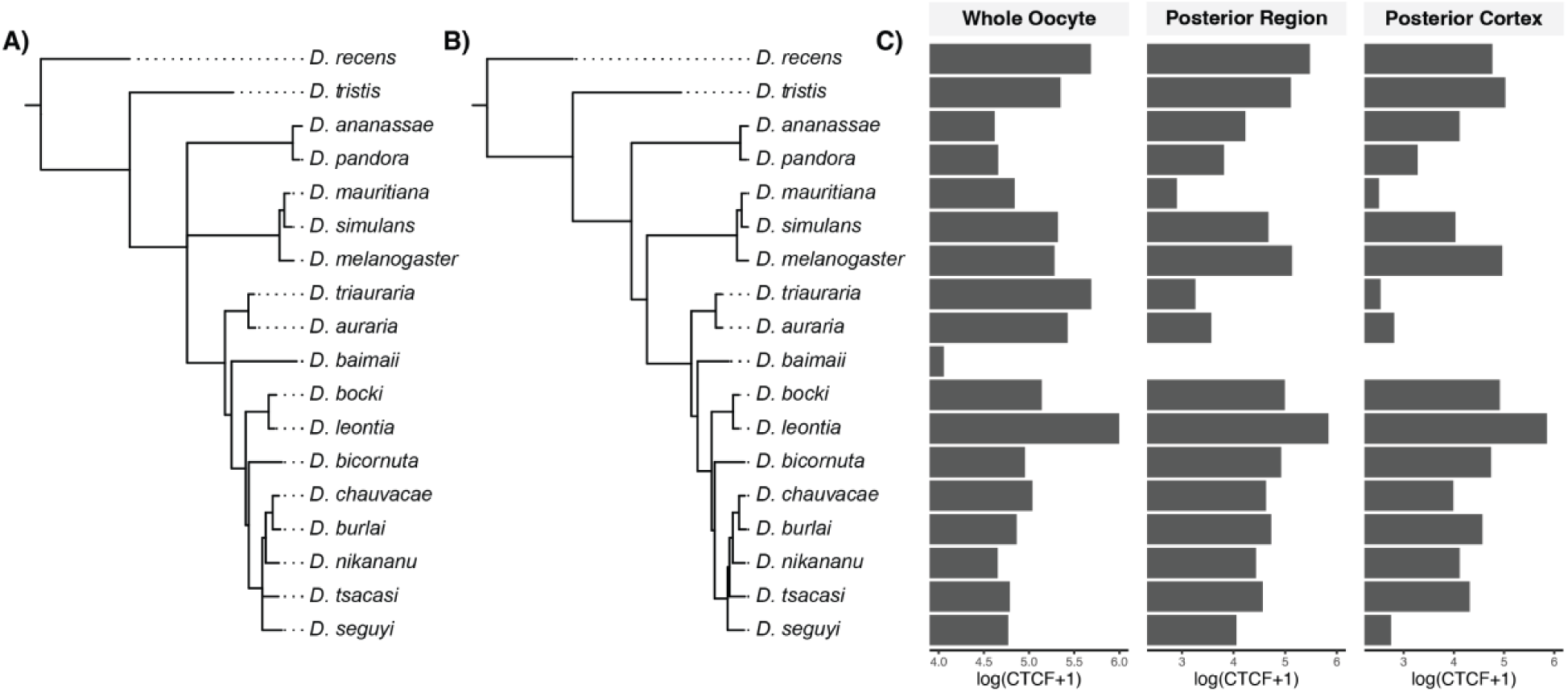
(A) Estimated Bayesian phylogram of host species based on 20 nuclear loci. All nodes have posterior probabilities >0.95. (B) Bayesian phylogram estimated using similar methods, but with a constrained topology to account for previously resolved relationships among subgroups within the *melanogaster* species group ((*montium*, *melanogaster*) *ananassae*) based on Turelli et al., (2018) and Suvorov et al., (2022). (C) Mean estimates of *Wolbachia* abundance (log-transformed CTCF) in the whole oocyte, posterior region, and the posterior cortex. For *D. simulans*, the mean values include both *w*Ri- and *w*Ha-infected individuals.

**Supplemental Figure 11.**
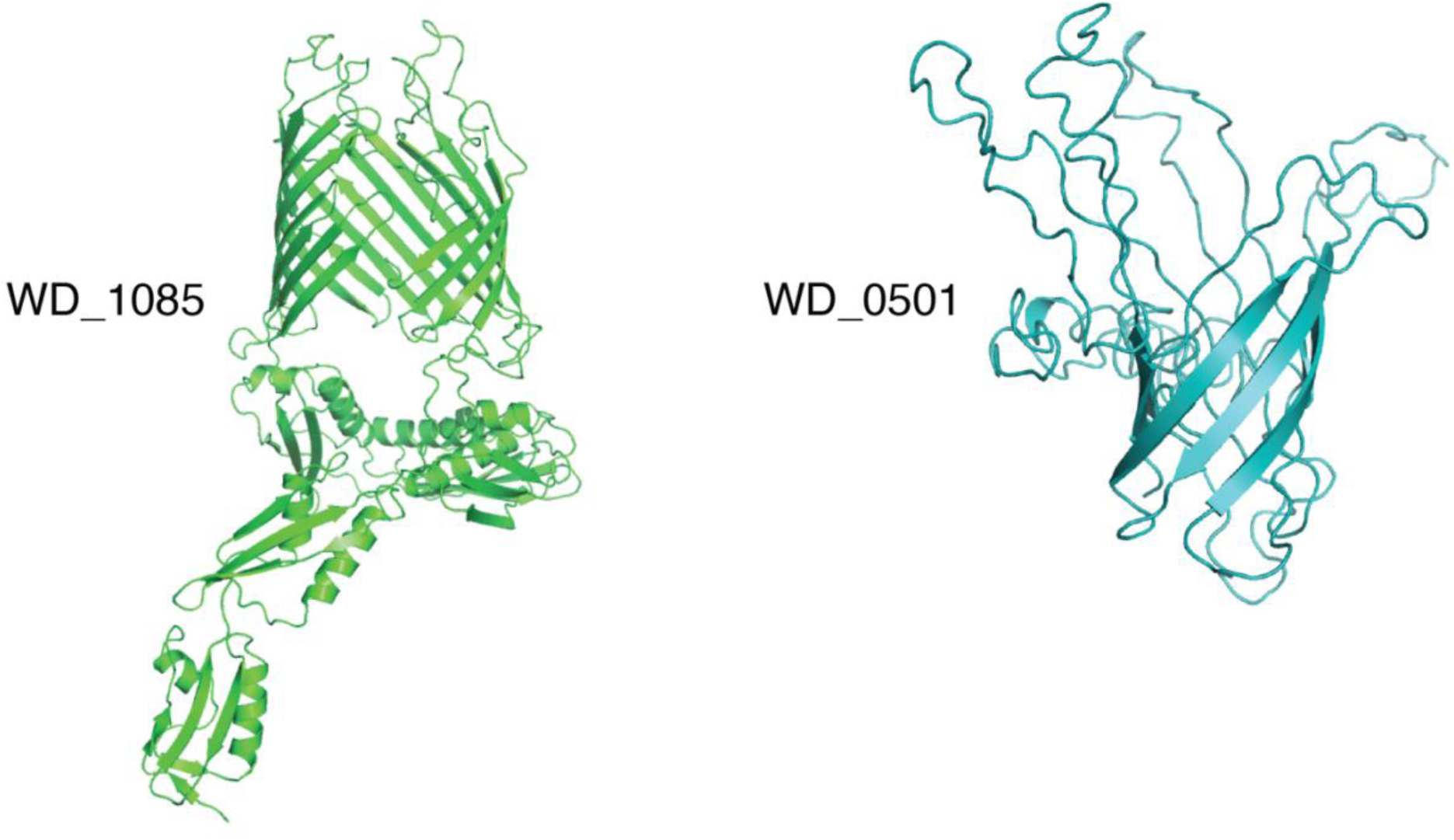
Illustrations of the homology models for the *Wolbachia* surface proteins WD_1085 and WD_0501.

**Supplemental Table 1.**
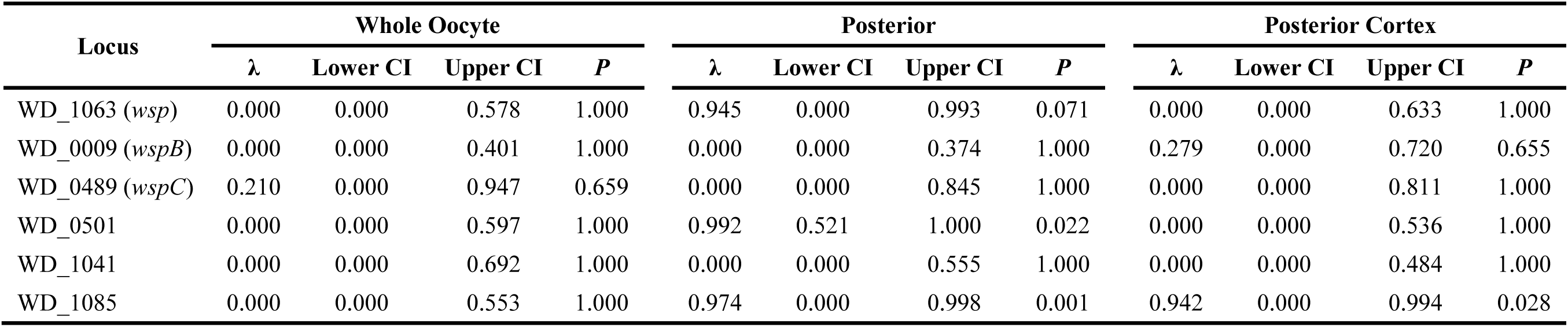
Estimates of phylogenetic signal (Pagel’s λ) with 95% confidence intervals (CI) for *Wolbachia* surface proteins.

**Supplemental Table 2.** Scaffold count, N50, and total assembly size of each new *Wolbachia* assembly.

| Host species | *Wolbachia* | Scaffold Count | N50 | Total Assembly Size |
| --- | --- | --- | --- | --- |
| *D. baimaii* | wBai | 209 | 9004 | 1135209 |
| *D. bicornuta* | wBic | 140 | 16267 | 1192611 |
| *D. bocki* | wBoc | 227 | 7115 | 1090928 |
| *D. burlai* | wBurl | 128 | 20905 | 1248003 |
| *D. chauvacae* | wCha | 406 | 4802 | 1300201 |
| *D. tsacasi* | wTsa | 59 | 29597 | 1132109 |
| *D. nikananu* | wNik | 83 | 22663 | 1257108 |
| *D. seguyi* | wSeg | 201 | 8880 | 1174927 |
| *D. tristis* | wTris | 114 | 20171 | 1264866 |
| *D. leontia* | wLeo | 186 | 9149 | 1096851 |

**Supplemental Table 3.** Summary of domain annotation results from HHPred for WD_1085 and WD_0501.

| Query name | Number | Hit | Name | Prob | E-value | Aligned cols | Target length | Query residues | Domain length |
| --- | --- | --- | --- | --- | --- | --- | --- | --- | --- |
| WD_1085 | 1 | COG4775 | BamA; Outer membrane protein assembly factor BamA [Cell wall/membrane/envelope biogenesis] | 100 | 3.10E-63 | 733 | 766 | 25-778 | 753 |
| WD_1085 | 2 | COG0729 | TamA; Outer membrane translocation and assembly module TamA [Cell wall/membrane/envelope biogenesis] | 100 | 2.00E-42 | 505 | 594 | 227-778 | 551 |
| WD_1085 | 3 | COG2831 | FhaC; Hemolysin activation/secretion protein [Intracellular trafficking, secretion, and vesicular transport] | 100 | 2.90E-39 | 490 | 554 | 237-778 | 541 |
| WD_1085 | 4 | KOG2602 | Predicted cell surface protein homologous to bacterial outer membrane proteins [General function prediction only] | 100 | 1.00E-31 | 433 | 457 | 315-778 | 463 |
| WD_1085 | 5 | PF19143.3 | Omp85_2 ; OMP85 superfamily | 99.94 | 2.60E-23 | 308 | 360 | 419-778 | 359 |
| WD_1085 | 6 | COG4775 | BamA; Outer membrane protein assembly factor BamA [Cell wall/membrane/envelope biogenesis] | 99.93 | 6.20E-21 | 569 | 766 | 25-637 | 612 |
| WD_1085 | 7 | COG4775 | BamA; Outer membrane protein assembly factor BamA [Cell wall/membrane/envelope biogenesis] | 99.92 | 2.80E-22 | 310 | 766 | 93-433 | 340 |
| WD_1085 | 8 | PF01103.26 | Omp85 ; Omp85 superfamily domain | 99.92 | 5.40E-22 | 306 | 424 | 454-778 | 324 |
| WD_1085 | 9 | PF03865.16 | ShlB ; Haemolysin secretion/activation protein ShlB/FhaC/HecB | 99.87 | 2.10E-19 | 285 | 310 | 416-742 | 326 |
| WD_1085 | 10 | COG4775 | BamA; Outer membrane protein assembly factor BamA [Cell wall/membrane/envelope biogenesis] | 99.61 | 2.50E-13 | 232 | 766 | 24-275 | 251 |
| WD_0501 | 1 | PF05736.14 | OprF ; OprF membrane domain | 99.78 | 3.00E-16 | 134 | 186 | 159-303 | 144 |
| WD_0501 | 2 | COG3637 | LomR; Opacity protein and related surface antigens [Cell wall/membrane/envelope biogenesis] | 99.75 | 1.20E-15 | 135 | 199 | 166-303 | 137 |
| WD_0501 | 3 | COG3047 | OmpW; Outer membrane protein W [Cell wall/membrane/envelope biogenesis] | 99.74 | 1.80E-15 | 144 | 213 | 159-303 | 144 |
| WD_0501 | 4 | PF06316.14 | Ail_Lom ; Enterobacterial Ail/Lom protein | 99.7 | 1.20E-14 | 179 | 199 | 3-303 | 300 |
| WD_0501 | 5 | PF19573.2 | DUF6089 ; Domain of unknown function (DUF6089) | 99.67 | 3.60E-14 | 186 | 226 | 4-303 | 299 |
| WD_0501 | 6 | PF13505.9 | OMP_b-brl ; Outer membrane protein beta-barrel domain | 99.65 | 8.40E-14 | 175 | 209 | 7-303 | 296 |
| WD_0501 | 7 | SCOP_d1p4ta_ | f.4.1.1 (A:) Outer membrane protein NspA {Neisseria meningitidis [TaxId: 487]} \| CLASS: Membrane and cell surface protei | 99.64 | 2.10E-13 | 154 | 155 | 95-303 | 208 |
| WD_0501 | 8 | PF01389.20 | OmpA_membrane ; OmpA-like transmembrane domain | 99.62 | 2.60E-13 | 173 | 202 | 95-303 | 208 |
| WD_0501 | 9 | SCOP_d1qjpa_ | f.4.1.1 (A:) Outer membrane protein A (OMPA) transmembrane domain {Escherichia coli [TaxId: 562]} \| CLASS: Membrane and | 99.61 | 4.60E-13 | 167 | 171 | 94-303 | 209 |
| WD_0501 | 10 | PF03922.17 | OmpW ; OmpW family | 99.6 | 3.40E-13 | 164 | 201 | 96-303 | 207 |

**Supplemental Table 4.** Summary of results for *Wolbachia* surface proteins.

| Gene | Description | Significant Phylogenetic Signal | Ectopic Expression |
| --- | --- | --- | --- |
| WD_1063<br>(wsp/RS04815) | surface antigen Wsp | None | Growth inhibition |
| WD_0009<br>(wspB/RS00060) | surface antigen Wsp paralog | None | Growth inhibition |
| WD_0489<br>(wspC/RS06475) | surface antigen, Wsp paralog | None | Lethality |
| WD_1085_(RS04910) | outer membrane protein assembly factor BamA | Oocyte posterior, posterior cortex | Growth inhibition and sensitive to microtubule checkpoints/inhibitors |
| WD_0501_(RS0225) | P44/Msp2 family outer membrane protein | Oocyte posterior | Slight growth inhibition |
| WD_1041_(RS04710) | peptidase M2 | None | Not tested |

